# A distinct growth physiology enhances bacterial growth under rapid nutrient fluctuations

**DOI:** 10.1101/2020.08.18.256529

**Authors:** Jen Nguyen, Vicente Fernandez, Sammy Pontrelli, Uwe Sauer, Martin Ackermann, Roman Stocker

## Abstract

It has been long known that bacteria coordinate their physiology with environmental nutrient, yet our current understanding offers little intuition for how bacteria respond to the second-to-minute scale fluctuations in nutrient concentration characteristic of many microbial habitats. To investigate the effects of rapid nutrient fluctuations on bacterial growth, we coupled custom microfluidics with single-cell microscopy to quantify the growth rate of *E. coli* experiencing 30 s to 60 min nutrient fluctuations. Compared to steady environments of equal average concentration, fluctuating environments reduced growth rate by up to 50%. However, measured reductions in growth rate were only 38% of the growth loss predicted from single nutrient shifts — an enhancement produced by the distinct growth response of cells grown in environments that fluctuate rather than shift once. We report an unexpected physiology adapted for growth in nutrient fluctuations and implicate nutrient timescale as a critical environmental parameter beyond nutrient concentration and source.

## Introduction

Our planet is sustained by the metabolic activities of microorganisms. In our gut, microbial communities break down nutrients into forms that we can take up and use; at sea, microbial growth affects the sequestration of carbon in the ocean and its release back into the atmosphere; and microbes in the soil convert organic molecules into forms that facilitate plant growth. These metabolic activities are often performed under conditions that depart from steady state. Rather, the quality and quantity of available nutrients often fluctuate rapidly due to microscale spatial heterogeneity, fluid flow, or host eating habits. Many host-associated or free-living microbes swim through resource landscapes that are highly heterogeneous at sub-millimeter scales (1; 2; 3) and thus experience rapid fluctuations in nutrient availability over seconds or minutes (4). Surface-attached microorganisms experience rapidly changing resources as a consequence of the movement of the liquid phase (5; 6; 7). To understand the impacts that microorganisms have on the physiology of their hosts and on global elemental cycles, we thus have to understand how individual bacteria respond to nutrient fluctuations. However, our understanding of microbial physiology draws heavily on knowledge derived from steady-state environments or single transitions between steady states (8; 9; 10). Microbial metabolism and growth under nutrient fluctuations remains a knowledge gap, largely due to the technical challenges of studying cells in highly dynamic environments. Here, we address this gap using single-cell growth experiments in a custom microfluidic device to show that rapid fluctuations substantially diminish growth, but also that bacteria can exhibit a fluctuation-adapted growth physiology that favors growth under frequent environmental change.

Recent advances in single-cell measurement techniques have laid foundations for considering the implications of second- and minute-scale fluctuations on bacterial growth and physiology. Single-cell measurements of bacterial mass at femtogram resolution have confirmed that individual bacteria add mass exponentially (11; 12). Experiments based on a groundbreaking microfluidic tool, the Mother Machine, revealed that the exponential growth of individual cells is stable over hundreds of generations (13), indicating that steady-state growth applies not only at the population level, but also to individuals. This seminal discovery has catalyzed major progress towards understanding the homeostatic regulation by which bacteria tune their growth and physiology to their environment, resulting for example in the Adder model of cell-size control (14; 15; 16) and hypotheses for its underlying mechanisms (17; 18; 19). This wealth of literature stems from and reinforces a long-standing paradigm: that each nutrient environment induces a characteristic steady-state growth rate (8; 9; 16), in which cells tightly regulate their size (16), proteome (20), and biosynthesis rates (9) in response to nutrient availability. The robustness of steady-state cell physiologies has led to growth laws that relate physiological traits, such as RNA–protein ratios (9; 21), with steady-state growth rate.

The expansive experimental and theoretical characterization of steady-state growth has led to its use as the framework to interpret bacterial physiology and ecology, even for dynamic environments. Currently, our understanding of bacterial responses to changes in the environment derives heavily from characterizations of physiological transitions from one steady state to another. Upon a nutrient shift out of steady state, cells initiate a cascade of responses that depend on the nutrient composition of the new environment (9; 10) and typically require hours to complete (22). Specific processes respond over distinct timescales — transcription over seconds, translation over minutes, cell division over hours (22) — and the progression of these physiological changes is reflected in a cell’s growth rate. The kinetics of growth transitions thus provide important insight into the strategies employed by bacteria and the ecological challenges under which these strategies have evolved (10; 23; 24).

While steady-state growth is highly informative when nutrients shift on timescales sufficiently long for cells to reach steady state before the next environmental change, it is unclear whether it provides an appropriate framework for understanding physiology when nutrients fluctuate on timescales of seconds or minutes. Under a steady-state framework, bacteria sense the availability of nutrient and induce the expression of the appropriate genes to transition their physiology toward the steady state characteristic of the instantaneous environment. However, the physiological ramifications of continuously tracking an environment that switches faster than the time required for complete steady-state adaptation are unclear. For example, alternating regulation between two steady states may produce cells with proteins that are typically not co-expressed. Proteins expressed during prior exposures to a certain condition might reduce lag times when that condition returns (25; 26), yet unnecessary gene expression can also reduce growth rate (20). How these slower gene expression-based responses might integrate to affect bacterial growth in rapid nutrient fluctuations is unclear due to the lack of systematic studies of single-cell growth in dynamic nutrient conditions at fast timescales.

In this study, we characterized the rate and kinetics of bacterial growth under fluctuations between two fixed nutrient concentrations on timescales of seconds to minutes. Using a custom microfluidic device that precisely controls the time dependence of nutrient delivery, we quantified the growth dynamics of thousands of individual *E. coli* cells exposed to identical, periodic nutrient fluctuations with periods as short as 30 s. We found that nutrient fluctuations reduce growth rate by up to 50% compared to a steady nutrient delivery of equal average concentration. However, the measured loss is considerably (38%) smaller than the growth loss expected from a null model based on the measured growth response to a single shift in nutrient concentration. We propose and provide evidence for a new growth physiology that is markedly distinct from steady-state physiologies and alleviates growth loss in fluctuating nutrient environments. Thus, this work implicates temporal variability as a fundamental parameter for understanding bacterial physiology in dynamic habitats.

## Results and Discussion

### Exposing single bacteria to precisely controlled, rapid nutrient fluctuations

To determine how rapid nutrient fluctuations affect bacterial growth, we engineered a microfluidic device to rapidly switch between the delivery of two different nutrient concentrations (one labeled with fluorescein) while simultaneously imaging individual bacteria with time-lapse phase-contrast microscopy (**Fig. 1a**; **Methods**; **Supplementary Fig. 1**). We subjected surface-attached *E. coli* to square-wave nutrient oscillations with periods of 30 s, 5 min, 15 min or 60 min (**Fig. 1b**). Each experiment began by flowing cells from a growing batch culture into the device and allowing them to settle onto its lower glass surface for 10–15 min prior to initiating nutrient fluctuations (**Methods**; **Supplementary Fig. 2**). Switches between the two nutrient concentrations occurred while maintaining a constant flow rate and each switch completed in less than 3 s (**Supplementary Fig. 3**). The resulting nutrient signal was reliably experienced by the cells as a square-wave of equal time (half the period) in each concentration (**Fig. 1b**), with sharp transitions between concentrations (**Fig. 1c**). Due to the high flow rates and channel depth (60 μm) used (**Supplementary Table 1**), the composition of the nutrient media was not altered by nutrient depletion or metabolite accumulation (**Supplementary Fig. 4**). Usually, one of the two cells emerging from each division was transported away by the flow (**Fig. 1d**), allowing us to acquire time series of thousands of individual cells (4,000–20,000) for each experiment. We confirmed that growth rates were independent of a cell’s position along the 10-mm-long region imaged within the microchannel (**Supplementary Fig. 4**) and therefore that cells experienced identical nutrient time series.

**Fig. 1:**
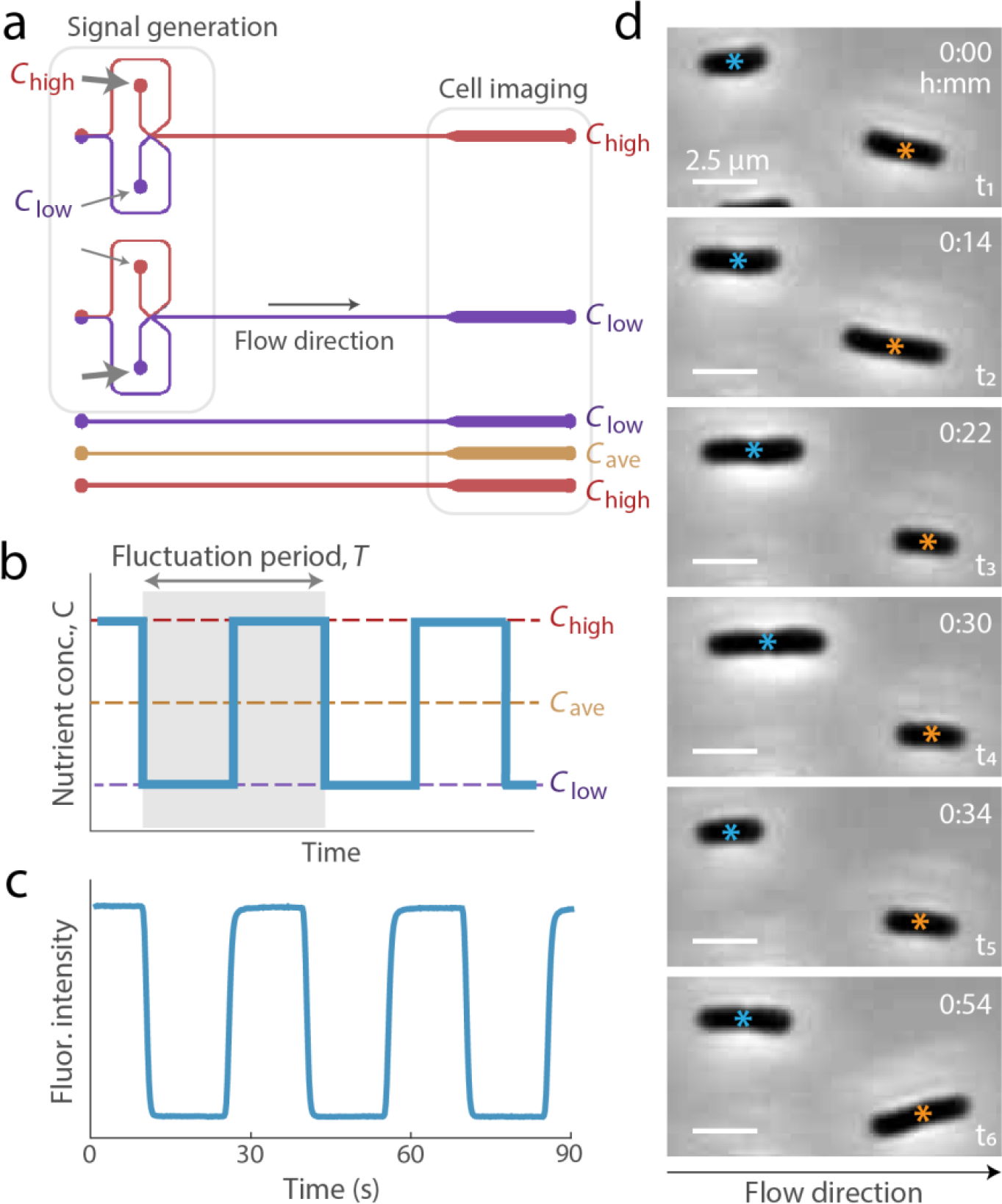
The Microfluidic Signal Generator (MSG) creates automated, precise high-frequency fluctuations in nutrient concentration while enabling single-cell microscopy. **a** Two channel configurations: the MSG for switching between two media (top) and straight channels for steady delivery of a single medium (bottom). The upstream portion of the MSG facilitates switching between media delivered to downstream cells via automated control over the pressure differences driving two nutrient media while maintaining a constant flow rate into the device. The wider downstream section fits over 10 imaging fields of view at 60x magnification. Each microfluidic device contains four channels: one MSG and three straight. The two MSG channels displayed here schematically represent the flow conditions that expose either *C*_low_ and *C*_high_ to the cells. **b** Bacteria were exposed to fluctuating signals in the form of even oscillations between a low and a high LB concentration (*C*_low_ and *C*_high_), with period, *T*, between 30 s and 60 min. Three control environments, *C*_low_, *C*_ave_, and *C*_high_, were run simultaneously with each fluctuating environment. **c** Fluorescein intensity illustrates the signal received at the cell imaging region over multiple oscillations (*T* = 30 s). Transitions between media are completed in less than 3 s. **d** Individual *E. coli* cells growing within the MSG. Cells were imaged at 117 sec intervals; timestamps of selected images are displayed in minutes. Cells divide between t_2_ and t_3_ (orange) and between t_4_ and t_5_ (blue), and can be seen elongating between other frames. One of the two cells emerging from division is typically swept away with the flow. Scale bars indicate 2.5 μm.

To isolate the role of nutrient fluctuation timescale from that of nutrient concentration, we switched between the same two nutrient concentrations for all fluctuating environments, a “high” and a “low” concentration of a complex growth medium (*C*_high_ = 2% LB, *C*_low_ = 0.1% LB), empirically chosen to avoid the saturation of growth rate (**Supplementary Fig. 5**). Three control experiments with steady nutrient concentrations were run in parallel with each fluctuating experiment: one at *C*_high_, one at *C*_low_, and one at *C*_ave_ = (*C*_high_+*C*_low_)/2 (*i.e*., 1.05% LB) (**Fig. 1a**). Importantly, the *C*_ave_ control provided cells the identical average and total nutrient as the fluctuating environments. Together, these steady controls enabled us to distinguish the effects of fluctuation timescale from those of nutrient concentration by providing reference growth rates in steady high, average and low nutrient concentrations, respectively (**Supplementary Fig. 5**).

### Growth rate rapidly responds to nutrient fluctuations

This microfluidic system allowed us to measure the growth rate of individual cells in steady and fluctuating nutrient concentrations with high precision and high temporal resolution. Growth rate, defined here as the rate at which volume doubles, was quantified from changes in cell volume between phase-contrast images of single cells acquired approximately every 2 min (**Methods**). For each cell imaged, we extracted the length and width using image analysis, and quantified cell volume, *V*(*t*), by approximating the cell as a cylinder with hemispherical caps (17; 27). Using the resulting time series of cell volume, *V*(*t*) (**Fig. 2a**), we computed the instantaneous single-cell growth rate, μ(*t*), from *V*(*t*+Δ*t*) = *V*(*t*) · 2^μΔ*t*^ (**Methods**). Growth rates in the steady controls stabilized within 3 h of the start of the experiments (**Supplementary Fig. 5**), so single-cell growth rates measured after 3 h were used to compute the steady-state growth rates *G*_high_, *G*_ave_ and *G*_low_, averaged across all single cells in *C*_high_, *C*_ave_ and *C*_low_, respectively (**Fig. 2b**). We confirmed with metabolomic profiling that the different steady-state growth rates between the three steady conditions resulted from proportional changes in nutrient uptake rates, rather than changes in the uptake of preferred metabolites (**Methods**; **Supplementary Fig. 6**).

**Fig. 2:**
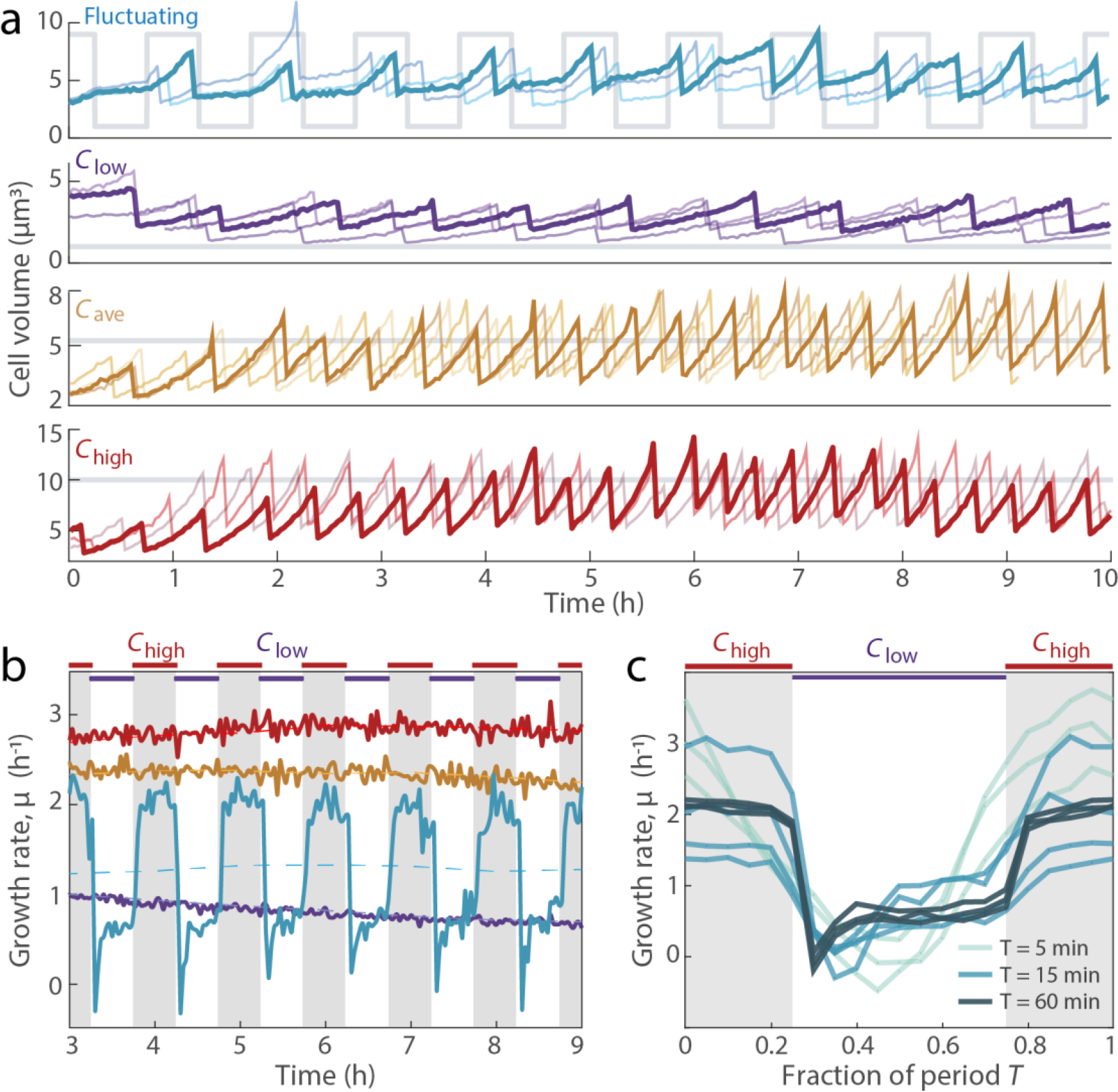
Nutrient upshifts and downshifts are followed by rapid adjustments in growth rate. **a** Single-cell volume trajectories from a fluctuating environment (fluctuation period *T* = 60 min) and control environments: steady *C*_low_, *C*_ave_, and *C*_high_. Each colored line tracks the repeated growth and division events of a single cell. Gray lines show the nutrient signal of each environment. **b** Instantaneous growth rate, μ, over time in steady and fluctuating environments. In the steady environments, μ is stabilized at steady-state growth by *t* = 3 h. In a fluctuating environment (*T* = 60 min), μ fluctuates with the nutrient signal. Each curve is a time-average of all instantaneous single-cell growth rates from each 2-min time bin, based on estimates of μ from at least 1842 cells per replicate experiment. Given noise in μ at the single-cell level (see **Supplementary Fig. 7**), the growth rate dynamics were best visualized by averaging many cells. **c** Instantaneous growth rate, μ, averaged across all single-cell growth rates as a function of the nutrient phase in fluctuating environments. Regardless of the timescale of nutrient fluctuation (*T*), μ is higher in *C*_high_ and lower in *C*_low_. Regions shaded in gray correspond to *C*_high_ phases of the fluctuating nutrient signal. Curves represent replicate experiments.

The growth rate in fluctuating nutrient conditions changed rapidly in response to the fluctuations. Because instantaneous growth rate dynamics from single cells were noisy (**Supplementary Fig. 7**), we averaged the single-cell growth rates at each time point to obtain instantaneous growth rates, which displayed strong and sharp fluctuations, changing more than two-fold within minutes (**Fig. 2b**). Instantaneous growth rate fluctuated with the immediate nutrient concentration, such that higher growth rates were observed in the high-nutrient phase and lower growth rates in the low-nutrient phase (**Fig. 2b**). When averaging the instantaneous growth rates from each phase of the nutrient signal, periodic changes were observed when nutrient fluctuated with a period of 5 min, 15 min or 60 min (**Fig. 2c**), indicating that growth rate responded to nutrient shifts in less than 2.5 min. Changes in growth rate that slightly precede changes in the nutrient signal are not an anticipatory response, but rather are caused by limits in the time-resolution of our measurements (**Supplementary Fig. 8a**). Similarly, fluctuations in growth rate were not resolvable with 30 s fluctuations, owing to the temporal resolution of image acquisition (2 min). Nevertheless, these results establish that rapid fluctuations in nutrient concentration lead to minute-scale fluctuations in instantaneous growth rate. We next asked how these nutrient fluctuations – and the resulting fluctuations in instantaneous growth rate – impact the mean rate at which individual cells grow.

### Rapid nutrient fluctuations reduce growth rate relative to steady conditions

The mean growth rate in fluctuating environments, *G*_fluc_, was consistently lower than the growth rate in the steady average conditions, *G*_ave_. For each fluctuating timescale, we computed *G*_fluc_ as the average of all instantaneous growth rates for all cells, measured from 3 h to the end of the experiment. For nutrient fluctuations of 30 s, 5 min, 15 min and 60 min periods, this measure yielded values of *G*_fluc_ of 1.93 ± 0.16 h^-1^, 1.53 ± 0.20 h^-1^, 1.15 ± 0.28 h^-1^ and 1.15 ± 0.13 h^-1^, respectively (mean ± standard deviation; *n* = 3–4 replicate experiments per condition, each with at least 1842 cells; **Fig. 3a**; **Supplementary Table 2**). The corresponding value of *G*_ave_ was 2.31 ± 0.18 h^-1^ (*n* = 13 replicate experiments; **Supplementary Table 2**). Accordingly, *G*_fluc_ was lower than *G*_ave_ by 16.5–50.2% of *G*_ave_.

**Fig. 3:**
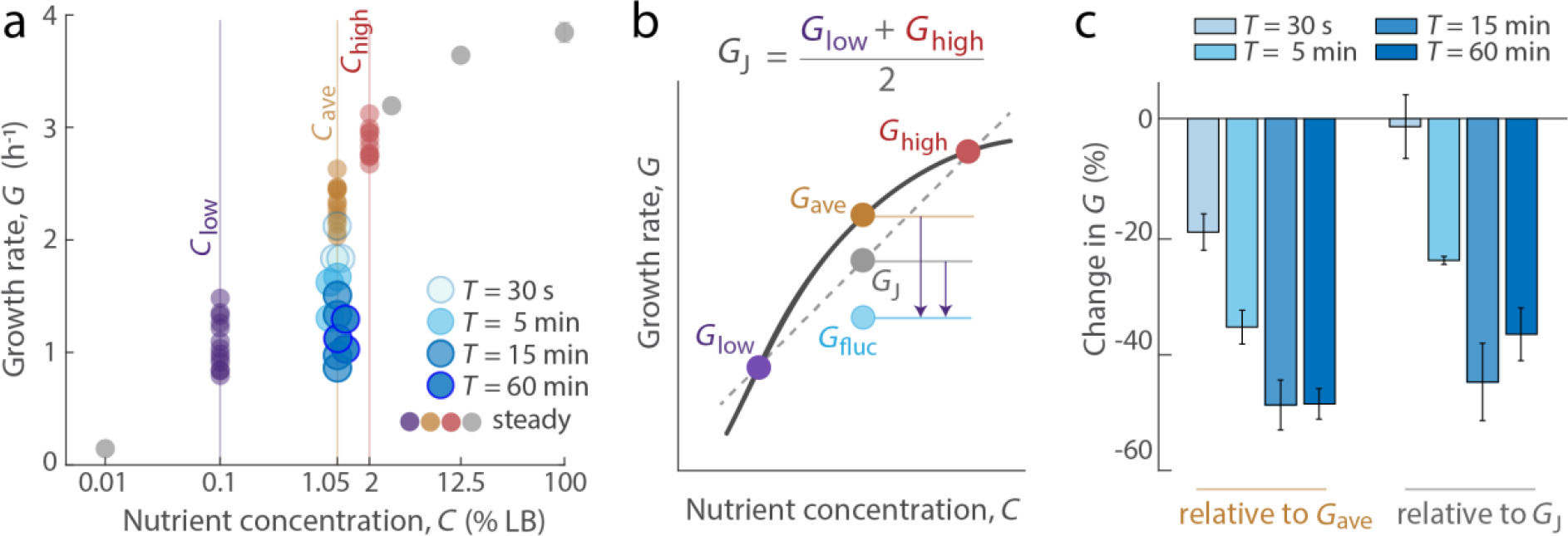
Rapid nutrient fluctuations reduce growth rate compared to environments of equal average nutrient concentration. **a** Cells in fluctuating environments of various period lengths (*T*) experienced the same time-averaged nutrient concentration as *C*_ave_ but grew at lower growth rates. Each point represents the mean growth rate of all individual cells measured from all time steps after the initial 3 h of each experiment. Colored points are replicate experiments for steady *C*_low_, *C*_ave_ or *C*_high_ or fluctuating nutrient conditions; gray points are additional steady nutrient concentrations that span nearly no growth to saturated growth. Error bars denote the standard error of the mean and are smaller than data points when not visible. **b** Schematic representation of Jensen’s inequality (*G*_ave_ > *G*_J_) and the relationship between *G*_ave_, the growth rate at steady nutrient *C*_ave_ = (*C*_low_ + *C*_high_)/2, and *G*_J_, the maximum growth rate expected for cells spending equal time at *C*_low_ and *C*_high_. *G*_fluc_ in our experiments is below even *G*_J_, indicating that the non-linear relationship between growth rate and nutrient concentration is not sufficient to explain why fluctuations decrease growth rate. **c** Percent change in growth rate for *G*_fluc_ at different period lengths relative to a steady-state reference, either *G*_ave_ or to *G*_J_. Change was calculated as (*G*_fluc_ - *G*_ave_)/ *G*_ave_ ×100 or (*G*_fluc_ - *G*_J_)/ *G*_J_ ×100 from conditions performed on the same day. Error bars denote the standard error of the mean.

This reduction in mean growth rate has a strong impact on bacterial population dynamics. For example, for an initial biomass of *M*_0_ = 1 μm^3^, the values of *G*_fluc_ for 30 s fluctuations (1.93 ± 0.16 h^-1^) and of *G*_ave_ (2.31 ± 0.18 h^-1^) correspond to a daily (*t* = 1 d) biovolume produced of 9 × 10^13^ μm^3^ and 5 × 10^16^ μm^3^, respectively (**Supplementary Table 3**). That is, two cells of equal initial volume, both growing exponentially (*M*(*t*) = *M*_0_ · 2^*Gt*^) with one at rate *G*_fluc_ and one at *G*_ave_, will differ in biomass production by over 500-fold in a single day, implicating the timescale of nutrient availability as an important parameter in the dynamics of bacterial populations.

Why do fluctuations reduce growth rate relative to the steady-state growth rate, *G*_ave_? The concave Monod curve offers a potential, purely mathematical explanation: steady-state growth rate increases less than linearly with nutrient concentration (**Fig. 3b**). This is an example of Jensen’s inequality, which states that for a concave function, the mean of the function (*i.e*., *G*_J_ = (*G*_low_ + *G*_high_)/2) is smaller than the function of the mean (*i.e*., *G*_ave_). Thus, Jensen’s inequality predicts from the steady-state growth function that fluctuations between *C*_high_ and *C*_low_ would result in a growth rate lower than *G*_ave_. Indeed, the mean growth rate based on Jensen’s inequality is *G*_J_ = 1.97 ± 0.16 h^-1^ (**Fig. 3b**), which is lower than *G*_ave_ = 2.31 ± 0.18 h^-1^. However, Jensen’s inequality does not explain the observed growth reductions (**Fig. 3c**): the growth rates from fluctuating environments, *G*_fluc_, were lower than *G*_J_ for all fluctuation periods except 30 s (**Supplementary Fig. 9**). Thus, the non-linear relationship between growth rate and nutrient concentration is not sufficient to explain the magnitude of the growth rate reduction caused by fluctuations.

We hypothesized that the reduction in growth rate results from the time required for cells to adopt the steady-state physiology characteristic of the current nutrient condition after each fluctuation. The prevailing paradigm for growth transitions presumes that cells initiate a physiological transition (mediated by differential gene expression) to the immediate nutrient environment, regardless of the nutrient timescale. In response to a shift in nutrient concentration (*e.g*., from *C*_low_ to *C*_high_), cells grow at rates lower than *G*_high_ for a period of several hours until the physiological transition is complete (**Fig. 4a**). The hours-scale transition in growth rate is a characteristic response to environments in which the nutrient condition shifts only once (10; 22; 23; 24). Based on this, growth dynamics are predictable from steady states when nutrient fluctuates on timescales sufficiently long for cells to complete their physiological transition before the environment again changes (28). When nutrients fluctuate on timescales faster than the time required to complete a physiological transition, the paradigm predicts that cells would be continuously transitioning and thus continuously growing at rates lower than steady state, causing *G*_fluc_ to be lower than *G*_J_.

**Fig. 4:**
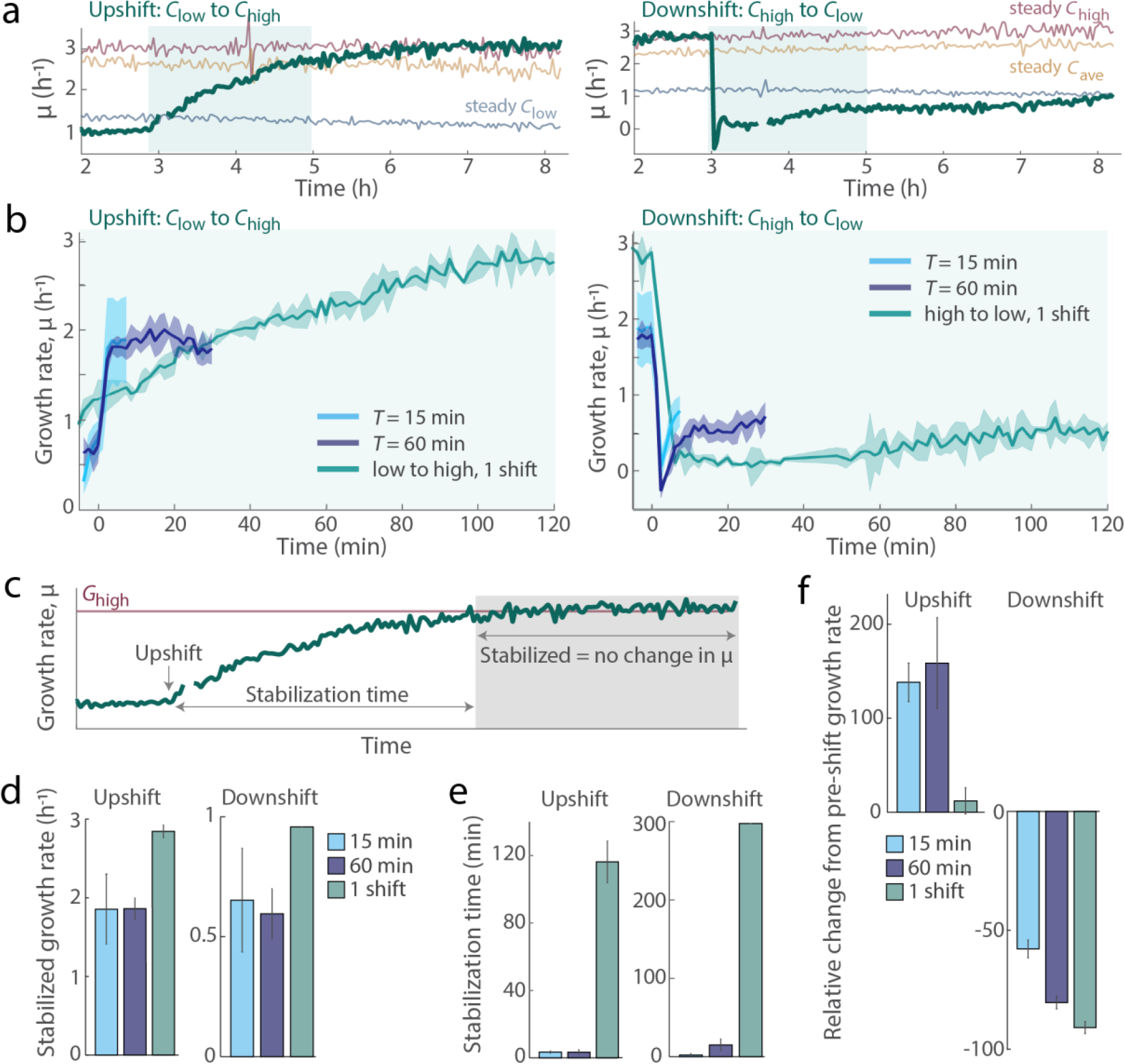
Growth rate responses differ between repeated fluctuations and single nutrient shifts. **a** Average growth rate of single cells over time in four conditions: a single shift in nutrient concentration (occurring at 3 h, after cells had reached steady-state growth in the initial condition), steady *C*_low_, steady *C*_ave_ and steady *C*_high_. On the left, a single nutrient upshift (shift from *C*_low_ to *C*_high_) and on the right, a single nutrient downshift (from *C*_high_ to *C*_low_). After each shift, growth rate gradually reaches steady-state growth in the post-shift condition. The growth rates in *C*_low_ before the upshift and in the steady *C*_low_ condition are both within the range of measured steady-state *G*_low_ (**Supplementary Table 2**). Data is from one representative experiment. The shaded region marks the portion of single-shift data plotted in **b**, which does not include the full progression to steady-state growth after a single up- or down-shift. **b** Growth rate from single cells in fluctuating nutrient conditions stabilize more rapidly and at a lower value when compared to the growth rate dynamics of cells experiencing a single shift. Data was aligned such that the nutrient shift in all conditions occurs at *t* = 0. Post-shift data in fluctuating environments are plotted up until the next shift occurs. Shaded error bars denote the standard deviation among replicate experiments (*n* = 3–4 for fluctuating conditions; n = 2 for single-shift conditions). **c** Growth rate is considered stabilized once the slope of the growth signal within a shrinking window reaches zero. Stabilization time is defined as the time between the nutrient shift and the time at which growth rate is stabilized. **d** The growth rate of fluctuation-grown cells stabilized at rates lower than steady-state *G*_high_ or *G*_low_. Cells experiencing 15 min and 60 min fluctuations stabilized at 1.86 ± 0.47 h^-1^ and 1.86 ± 0.13 h^-1^, respectively, after an upshift and at 0.65 ± 0.22 h^-1^ and 0.60 ± 0.10 h^-1^ after a downshift. Cells shifted once from steady state stabilized only upon reaching steady state *G*_high_ (2.84 ± 0.08 h^-1^) after an upshift or *G*_low_ (0.96 h^-1^) when growth rate stabilized after a downshift (only one of two replicates stabilized at *G*_low_ after 5h of post-shift observation). **e** Cells grown in fluctuations stabilize in growth rate within 3.8 ± 0.0 (*T* = 15 min) or 3.3 ± 1.4 min (*T* = 60 min) of each upshift and within 2.2 ± 1.9 min (*T* = 15 min) or 15.0 ± 7.6 min (*T* = 60 min) of each downshift (*n* = 3–4). Cells grown in steady environments stabilize hours after a single shift, 116.3 ± 12.4 min in the case of upshifts (*n* = 2) and at least 297.5 min after a downshift (one of two replicates stabilized after 5 h). **f** The initial change in growth rate in the minutes following a nutrient shift was greater in the cells experiencing fluctuations than in cells experiencing only a single shift. Over the first 7.5 min after each upshift, cells experiencing fluctuations increased in growth rate relative to their pre-shift growth rate (*t* = 0) by 138.3 ± 20.6% (15 min) or 158.7 ± 48.4% (60 min), compared to a 12.0 ± 14.0% increase in the single shift case. Over the 7.5 min after each downshift, cells experiencing fluctuations decreased in growth rate by 57.8 ± 3.7% (15 min) or 80.4 ± 2.6% (60 min), compared to a 90.9 ± 2.5% decrease in growth rate in cells after a single shift. Error bars denote the standard deviation between replicates (*n* = 2–4).

### Growth rate responses differ between repeated fluctuations and single nutrient shifts

To determine whether the time spent physiologically transitioning can explain the reduction in *G*_fluc_ from *G*_J_, we compared the growth rate dynamics between cells exposed to fluctuations and cells exposed to a single up- or down-shift in nutrient concentration after having grown in steady conditions. In single-upshift experiments, cells growing steadily at *G*_low_ were switched to *C*_high_, while in single-downshift experiments cells growing steadily at *G*_high_ were switched to *C*_low_. These single-shift experiments enabled us to quantify the growth rate response as cells transitioned between two steady states, including the time for the growth rate adjustment to complete (**Fig. 4a**). In single-upshift experiments, growth rate gradually increased from *G*_low_ until it reached the steady-state value *G*_high_ after 2–3 h. In single-downshift experiments, growth rate dropped sharply from *G*_high_ down to 10% of *G*_low_, then gradually increased until it reached the steady-state value *G*_low_ after 5 h (**Fig. 4a**).

In contrast to cells experiencing single shifts in nutrient concentration, cells grown in fluctuations did not reach *G*_high_ or *G*_low_ as target set points. Instead, growth rate in fluctuations stabilized at values lower than steady-state values. To accurately quantify instantaneous growth rate dynamics in fluctuating environments, we focused on the fluctuations of 15 and 60 min periods, which were better resolved with our 2-min imaging interval. In fluctuations, we did not observe the transition characteristic of single shifts (**Fig. 4b**). Instead, growth rate under fluctuations stabilized at 66% of *G*_high_ after each upshift and at 63–68% of *G*_low_ after each downshift (**Fig. 4c, d**; **Supplementary Table 4**), indicating that cells grown in fluctuations are not continuously transitioning towards *G*_high_ or *G*_low_ in response to their immediate environment. These lower stabilized growth rates resulted in time-averaged growth rates (*G*_fluc_) that were smaller than *G*_J_. These observations show that the effect of rapid nutrient fluctuations on growth cannot be understood by reference to a sequence of single up- and down-shifts, and do not support the hypothesis that the reduction in growth rate under fluctuations results from continuous physiological transitions toward growth at *G*_high_ and *G*_low_.

The stabilization of growth rate was considerably faster in rapidly fluctuating environments than in steady environments perturbed by a single nutrient shift, indicating that cells grown under fluctuations have a different growth physiology from cells growth under steady nutrient conditions. While growth rate adjustments required hours after a single shift, instantaneous growth rate in fluctuating conditions stabilized within minutes of a nutrient shift (**Fig. 4b**): 3–4 min after each nutrient upshift and 2–15 min after each downshift (**Fig. 4e**). This stabilization of growth rate on a timescale of minutes is more consistent with a stable growth physiology, *i.e*. a single cellular composition that grows at different rates due to changes in metabolite flux (**Fig. 4f**), rather than a physiology that is continuously in transition. Our results thus suggest the existence of a distinct physiology that cells adopt upon experiencing rapid nutrient fluctuations.

### Rapid nutrient fluctuations induce a novel growth physiology

To demonstrate that bacteria can adopt a distinct physiological state when exposed to rapid nutrient fluctuations, we performed an experiment in which cells growing under steady *C*_low_ for at least 3.5 h were then exposed to fluctuations with period *T* = 60 min. We found that the first nutrient upshift induced the gradual increase in growth rate characteristic of the physiological transition between *G*_low_ and *G*_high_ (**Fig. 5a**). Subsequent upshifts displayed faster growth rate adjustments that increasingly resembled that characteristic of cells grown in fluctuating conditions (**Fig. 5a**). This transition confirms that cells can adopt a fluctuation-induced growth physiology, induced by repeated nutrient shifts.

**Fig. 5:**
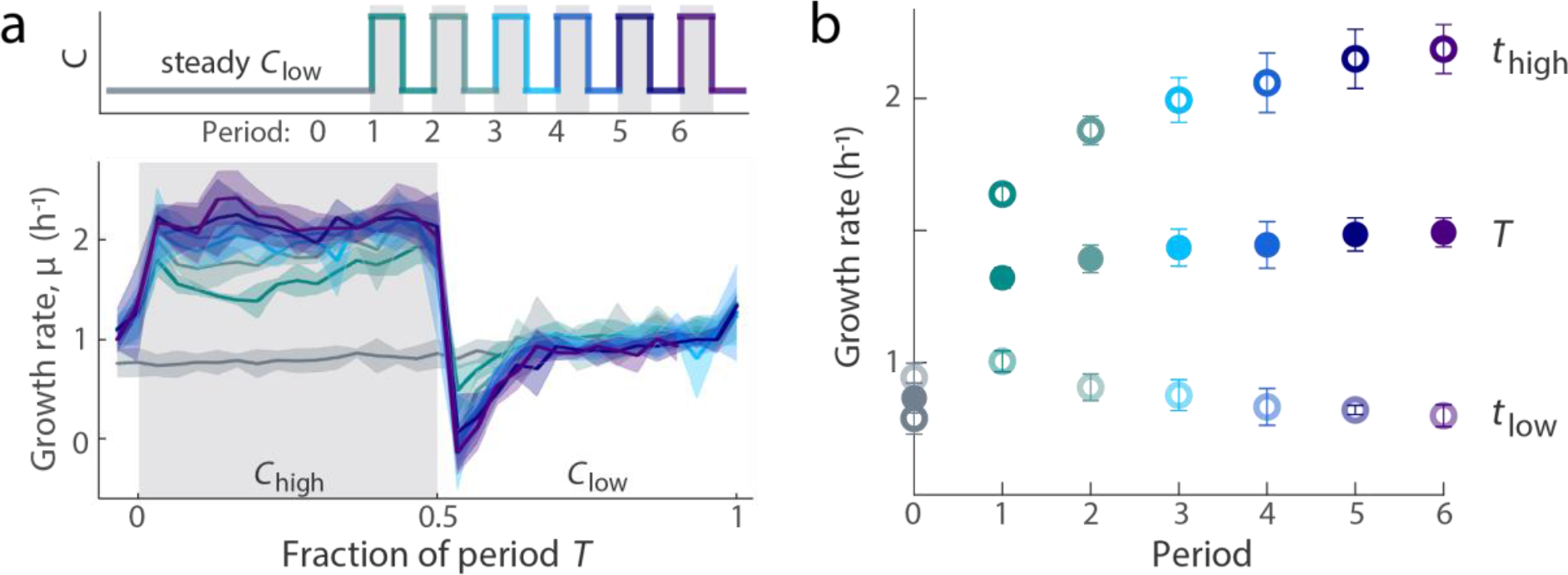
Rapid nutrient fluctuations induce the transition to a novel growth physiology. **a** After at least 3.5 h of growth in steady *C*_low_, cells experience the onset of nutrient fluctuations (*T* = 60 min) between *C*_low_ and *C*_high_. The growth rate dynamics in response to each successive nutrient period transition from the dynamics observed in response to a single shift to those observed in fluctuating environments (**Fig. 4b**). The growth rate dynamics of the successive periods, including the 60 min preceding the first nutrient upshift (“period 0”), are overlaid to show the stabilization of the growth rate signal. Shaded error bars denote the standard deviation among replicate experiments (*n* = 3). **b** Stabilization of growth rate dynamics with successive nutrient periods. The mean growth rate following each downshift (half period in *C*_low_, *t*_low_), each upshift (half period in *C*_high_, *t*_high_) and across the full period (*T*) adjusts and stabilizes by the third period. Error bars denote the standard error of the mean (*n* = 3) and are smaller than points when not visible.

The transition to a stable growth physiology occurred within 2–3 h of the onset of fluctuations. The growth following each successive nutrient upshift (growth in *C*_high_) increased during this 2–3 h transition time, offsetting the decreasing growth following each successive downshift (growth in *C*_low_) (**Fig. 5b**). In both nutrient concentrations, growth ceased to differ significantly from the third nutrient fluctuation period onwards (**Fig. 5b**). The overall increase from initial to stabilized growth suggests that the physiological transition to a fluctuation-induced physiology enhances growth in fluctuating environments. This enhancement is potentially a physiological trade-off, reducing growth rate in low nutrient concentration to increase the cell’s potential to quickly adjust to a higher growth rate when nutrient concentration becomes high (**Fig. 5b**). Indeed, in the minutes after a nutrient shift, the growth rates of cells in fluctuating conditions were higher than those of cells exposed to a single shift (**Fig. 4f**; **Supplementary Fig. 8b**). Thus, instead of continuously transitioning between steady-state physiologies, cells exposed to rapid nutrient fluctuations adopt a distinct, stable physiology that alleviates the challenges of growth in unsteady conditions.

### Growth in rapid nutrient fluctuations is higher than predicted by steady-state growth models

To quantify the advantages of a fluctuation-induced physiology, we compared mean growth rates under fluctuations with mean growth rates predicted by a null model based on the measured single-shift responses. The null model assumes the absence of a fluctuation-induced physiology, so that upon each shift in nutrient cells initiate the transition toward the corresponding physiological steady state, and uses the times series of the observed single-shift growth rate transitions (**Fig. 4a**) to predict the growth rate dynamics and thus *G*_fluc_ (**Fig. 6**). For long fluctuation periods (*T* = 12–96 h), each nutrient phase (duration *T*/2) is sufficiently long for cells to complete the 2–5 h physiological transition to a new steady state before the ensuing shift (**Supplementary Fig. 9**). As the period *T* increases, the null model thus predicts that the growth rate tends toward *G*_J_ (**Fig. 6**). As the period decreases, cells spend a larger fraction of time transitioning, causing a larger predicted reduction in growth rate relative to *G*_ave_ (for example, 17% below *G*_ave_ for *T* = 96 h, but 31% below *G*_ave_ for *T* = 12 h; **Supplementary Table 5**). In the absence of the fluctuation-induced physiology, the reduction in growth rate continues to increase as the period further decreases. The null model predicts values of *G*_fluc_ that are 44% below *G*_ave_ for *T* = 60 min and 50% below *G*_ave_ for *T* = 30 s (**Supplementary Table 5**). Because these nutrient periods are shorter than the physiological transition time, the null model assumed the cells were “low-adapted”: able to grow at *G*_low_ immediately upon experiencing *C*_low_ and, upon shifts to *C*_high_, displaying the growth rate dynamics of a single-shift toward growth at *G*_high_ (**Supplementary Fig. 9**). An alternative null model (see “high-adapted” in **Supplementary Fig. 9**) produced the same trend: faster fluctuations further decreased the predicted value of *G*_fluc_.

**Fig. 6:**
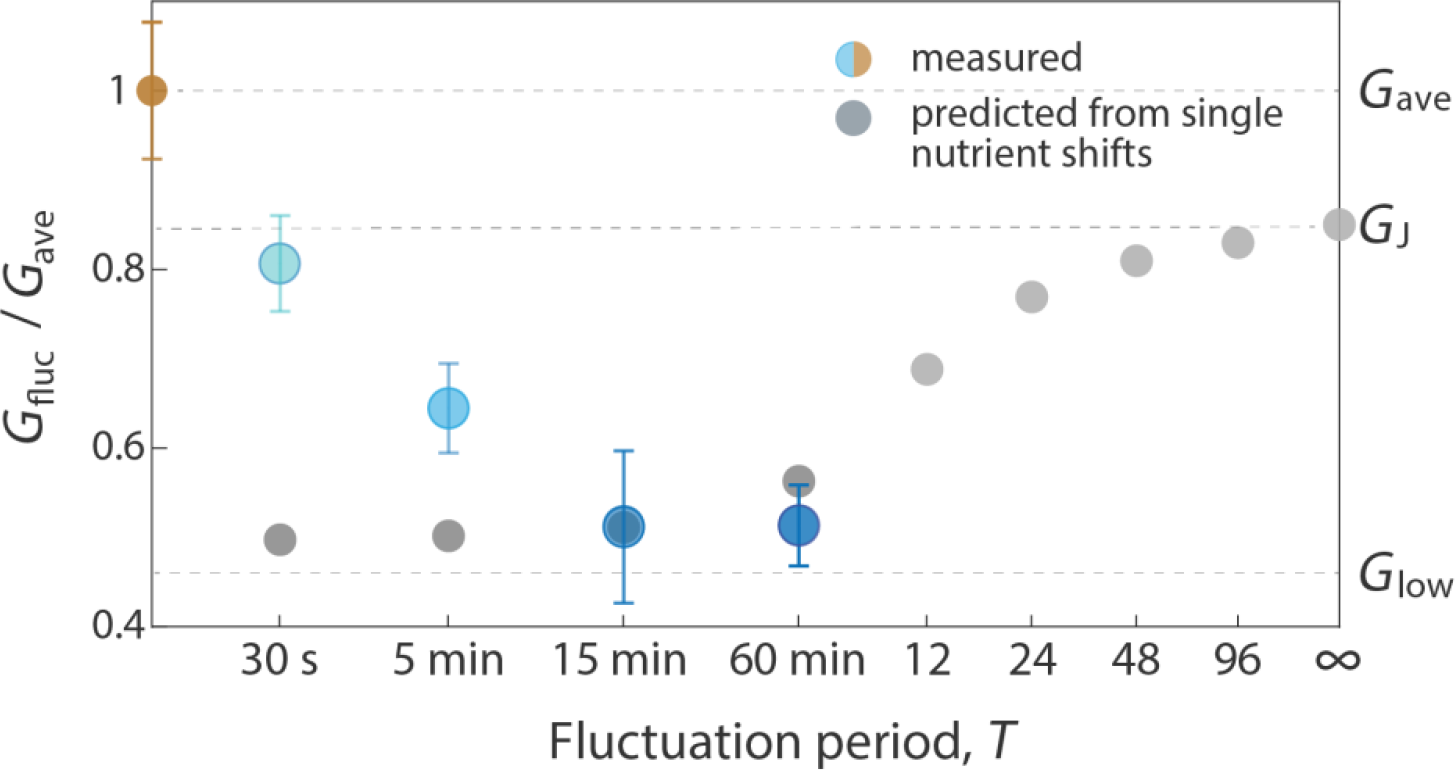
Growth rate under rapid nutrient fluctuations is sensitive to timescale and higher than predicted by steady-state models. Growth rates under rapid nutrient fluctuation (blue), expressed as a fraction of the growth rate in the steady average nutrient environment (*G*_ave_; yellow), are higher than predicted from data on single shifts between steady states. Each measured point represents the time-averaged growth rate *G* and standard deviation between replicates (*n* = 3–4). Predicted values of *G*_fluc_ (gray) reach a maximum of *G*_J_ when the fluctuating nutrient timescale is infinitely long relative to the time required to transition from growth at *G*_low_ to *G*_high_, and vice versa. At slow nutrient timescales (*T* = 12 h or more), predicted *G*_fluc_ was based on the growth rate dynamics measured from single nutrient up- and down-shifts. When each nutrient was shorter than the stabilization time (*T* = 60 min or less), the predictions considered cells were low-adapted and able to grow at *G*_low_ immediately upon experiencing *C*_low_. Thus growth rate during the *C*_high_ phases of the fluctuating signal followed the measured dynamics of the single upshift response between *t* = 0 and the length of the simulated *C*_high_ phase (**Supplementary Fig. 9**). Steady-state *G*_high_ is never reached, and growth returns immediately to *G*_low_ when the environment returns to *C*_low_. Alternative models to predict *G*_fluc_ under rapid nutrient timescales are presented in the Supplementary Information. All models predict lower *G*_fluc_ with decreasing period lengths (*T*) (**Supplementary Fig. 9; Supplementary Table 5**) — the opposite of the trend observed from the experimental values of *G*_fluc_. The deviation between measured and predicted *G*_fluc_ differentiates the growth physiology at rapid fluctuation timescales from the growth behaviors expected from steady state, and highlights the timescale-dependent nature of the growth advantage conferred by a physiology adapted for growth under rapid nutrient fluctuations.

This trend in the null model is the opposite of that displayed by the measured values of *G*_fluc_ for rapid fluctuations. With the observed fluctuation-induced growth rate dynamics, the experimentally observed value of *G*_fluc_ increases with decreasing fluctuation period: *G*_fluc_ is 50% below *G*_ave_ for *T* = 60 min, but only 16.5% below *G*_ave_ for *T* = 30 s (**Fig. 6**). Compared to the growth rates predicted from the null model, the experimental *G*_fluc_ values represent 21.9 ± 6.0% less growth loss relative to *G*_ave_ for *T* = 5 min and 38.2 ± 4.1% less growth loss for *T* = 30 s (**Fig. 6**). This increase in the measured value of *G*_fluc_ over that predicted by the null model represents the growth advantage in fluctuating conditions afforded by the fluctuation-adapted growth physiology. These results demonstrate that growth rate in rapid fluctuations is qualitatively and quantitatively distinct from steady-state growth dynamics.

An outstanding question concerns how cells sense that the environment is rapidly fluctuating and initiate the transition to a fluctuation-induced growth state. It is unclear how the fluctuation-induced physiology is achieved, in part because we do not know whether gene expression differs between cells growing in fluctuations and cells growing at steady state. Alterations in gene expression can be beneficial when environments change on extremely rapid timescales: for example, increased expression of photoprotection proteins has been demonstrated to increase the growth yield of plants exposed to minute-scale fluctuations in light (29). Alternatively, post-translational activation of “spare” ribosomes has been implicated as a bacterial strategy to increase growth rate rapidly upon nutrient upshift (30; 23; 24). The regulation of a fluctuation-induced physiology may of course combine gene expression and post-translational controls. Indeed, upon sensing depleted intracellular amino acid levels, *E. coli* has been observed to induce broad responses to increase energy production, ribosome levels and translational capacity (31). These observations have begun to probe the diversity of strategies that life has evolved to cope with the challenges of environmental change. By reporting a novel growth physiology in rapidly fluctuating environments, we contribute a new framework by which to pursue an understanding of bacterial growth that is relevant to realistic habitats.

## Conclusions

We found that when bacteria are exposed to nutrient fluctuations their growth rate is reduced compared to that observed in steady average nutrient conditions, even when nutrient fluctuated on timescales as rapid as seconds. These reductions are not explained by the current paradigm in bacterial physiology, which holds that cells experiencing a shift in their nutrient environment will transition to the steady state growth physiology of the post-shift environment. This paradigm was developed from experiments with environments that shift only once. Here, we report that rapidly fluctuating nutrient environments induce a novel bacterial physiology with growth responses fundamentally distinct single shifts from steady state that enables bacteria to grow faster in fluctuations than expected from the existing paradigm.

This work establishes nutrient timescale as a fundamental parameter characterizing bacterial environments and bacterial growth. In addition to chemical composition and temperature, we propose the temporal delivery of nutrients as a major determinant of growth, forming a third axis of nutrient environments to consider when studying bacterial behavior. The exposure of bacteria to rapid nutrient fluctuations is evident in *in situ* measurements (5; 32; 33), laboratory experiments (34; 35) and models (36). However, the physiological consequences of rapid nutrient fluctuations had not previously been investigated. Only through direct experiments designed to test the role of nutrient fluctuations in a controlled manner could nutrient timescale emerge as a key parameter for bacterial growth. By quantifying growth rate across a range of fluctuation timescales relevant to bacterial habitats, this work has uncovered a strong dependence of bacterial growth on the temporal dynamics of nutrient concentration, highlighting the importance of temporal variability (and by extension, microscale heterogeneity) when considering bacterial growth in realistic environments.

The fluctuation-induced growth physiology in *E. coli* opens the door to the discovery of novel forms of regulation by which bacteria coordinate their growth with nutrient availability. First, this study focuses on a single type of fluctuation – square waves – but huge complexity characterizes the time series of nutrient signals in the wild, including different fluctuation magnitudes, duty cycles, and degrees of randomness. Further studies with diverse patterns of nutrient fluctuations may yield additional strategies of bacterial growth in temporally variable environments. It is also possible that distinct strategies of growth in complex temporal environments have evolved between bacterial species that occupy distinct ecological niches. Taxon-specific specializations for the timescale of light fluctuations have long been observed in plants (37), and more recently in microbial response times to inputs of water to dry soils (38).

Our results illustrate how identical environmental shifts can induce different responses depending on the timescale at which the shifts are delivered, demonstrating the need to account for temporal variability in the environment at timescales that have been mostly ignored to date. Understanding the diversity of growth responses to realistic features of microbial environments will bring us closer to the establishment of general frameworks for bacterial growth in natural ecosystems and the discovery of mechanistic links between the interactions that occur on the scale of single cells, populations and communities.

## Methods

### Bacterial Strain

All experiments in this study were performed with the same *E. coli* strain, K-12 NCM3722 Δ*motA*. The background strain, K-12 NCM3722, has been sequenced (39) and physiologically characterized to be prototropic (40). The motility mutant (Δ*motA*) lacks flagella, facilitating long-term observation in microfluidics (16).

### Growth Media

#### Batch culture medium

MOPS medium (Teknova) supplemented with 0.2% glucose w/v and 1.32 mM K_2_HPO_4_ was used for overnight and seed batch cultures. All batch cultures contained 3 mL of supplemented MOPS medium inoculated with *E. coli*.

#### Microfluidics medium

Lysogeny broth (LB) composed of tryptone (10g/L), yeast extract (5 g/L) and NaCl (10g/L) was used for all microfluidic nutrient conditions. For the microfluidic experiments, the same stock solution of 100% LB was mixed with an equimolar NaCl solution (2.5g NaCl in 250 mL 0.22 μm filtered water (Millipore Millipak Express 40, catalog no. MPGP04001) to prepare three dilutions: low, average, and high. To avoid bubble formation within the microfluidics, the NaCl solution was freshly autoclaved the day of each experiment and then cooled before preparing the LB dilutions. The high LB (2%) mixed 2mL full LB into 98mL salt solution. The low LB (0.1%) mixed 5mL of the high LB solution with 95mL salt solution. Afterwards, the high LB solution was labeled with 0.26 nM sodium fluorescein, to allow visual calibration of switching between mediums. All solutions were adjusted to pH 7 with NaOH. Equal parts of low and high LB were mixed to produce the average LB control; hence the average LB medium contained 0.13 nM sodium fluorescein. This fluorescein addition had no effect on growth rate (**Supplementary Fig. 2a**). Furthermore, we confirmed with metabolomic profiling that the different steady-state growth rates between the three steady conditions resulted from proportional changes in nutrient uptake rates, rather than changes in the nutrients being metabolized (**Supplementary Fig. 6**). Growth media were loaded into plastic 10 mL syringes (Codan) or glass vials (VWR, cat. no. 548-0154) and warmed to 37 °C at least 3 h prior to the start of each experiment.

### Metabolomics characterization of growth media

To determine how diluting LB affected the consumption of its nutrient components, we collected the supernatant from 20 mL batch cultures grown at 37 °C shaking in 125 mL Erlenmeyer flasks. Twelve flasks were prepared in parallel, four of each nutrient concentration (*C*_low_, *C*_ave_, *C*_high_). Three of each concentration was inoculated with cells, while the fourth flask was kept bacteria-free and sampled as a blank control. The inoculum was prepared with the same overnight and seed culture preparation used for the microfluidics experiments (see **Cell Preparation**). Once the seed culture reached an OD_600_ of 0.1 (grown in MOPS medium with 0.2% glucose), nine 1 mL aliquots were centrifuged for 2 min at 2500 rcf (Eppendorf, Centrifuge 5424 R) and the MOPS-based supernatant removed. The cell pellet was gently resuspended in the final growth medium (*C*_low_, *C*_ave_ or *C*_high_) and then inoculated into the appropriate flask. Each flask was sampled every 30 min by centrifuging 500 μL of culture for 5 min at 2500 rcf. Then 100 μL of supernatant was removed from the top of each tube and stored in a 96-well plate (Thermo Scientific). Samples were kept on ice when in 1.5 mL microcentrifuge tubes (Sarstedt AG & Co.), and then at −20 °C when in the plate. The samples were thawed and diluted 1:10 in milliQ water prior to measurements with flow injection time-of-flight mass spectrometry (FIA-QTOF).

The different steady-state growth rates we measured arise from proportional changes in metabolite uptake, rather than shifts in the preferential uptake of some metabolites among the different concentrations of LB (*C*_low_, *C*_ave_, *C*_high_). To confirm that differences in growth resulted from different levels of nutrient uptake and not different consumed nutrient sources, depletion of extracellular metabolites was measured using flow injection time-of-flight mass spectrometry (FIA-QTOF) (**Supplementary Fig. 6**). Untargeted metabolomics measurements were performed with a binary LC pump (Agilent Technologies) and a MPS2 Autosampler (Gerstel) coupled to an Agilent 6520 time-of-flight mass spectrometer (Agilent Technologies) operated in negative mode, at 4 Ghz, high resolution, with a *m/z* (mass/charge) range of 50-1,000 as described previously (41). The mobile phase consisted of isopropanol:water (60:40, v/v) with 5 mM ammonium fluoride buffer at pH 9 at a flow rate of 150 μl/min. Raw data was processed and analyzed with Matlab (The Mathworks, Natick) as described in (Fuhrer et al., 2011) and ions were annotated against the KEGG database (restricted to *Escherichia coli*) with 0.003 Da tolerance. We detected 284 metabolites from batch cultures inoculated into *C*_ave_ or *C*_high_ (**Supplementary Table 6**) and monitored their depletion to find that all detectable metabolites displayed comparable dynamics in the two nutrient concentrations (**Supplementary Fig. 6**). This suggests that the different growth rates observed among nutrient concentrations arise from differences in nutrient flux, not differences in the composition of nutrient consumed. Metabolites in *C*_low_ were below the detection limit.

### Cell preparation

Cells for each experiment were grown in two batch cultures, the overnight culture and the seed culture, before entering the microchannels. The overnight culture was inoculated directly from a −80 °C glycerol stock into 3 mL of supplemented MOPS medium and shaken for 12-16 h at 37 °C at 200 rpm. The next morning, cells from the overnight culture were diluted to achieve 3 mL of supplemented MOPS medium with an initial OD_600_ below detection, generally a 1:1000 or 1:2000 dilution. This seed culture was used to inoculate microchannels once cells reached an OD_600_ between of 0.07 – 0.10.

### Microchannel fabrication

Microfluidic channels with a depth of 60 μm were cast in PDMS from a custom-made master mold (**Fig. 1a**) such that all four channels (1 Microfluidic Signal Generator (MSG) for fluctuating environments and 3 straight channels for steady environments) were present on the same device. Each PDMS device was bonded to a glass slide by plasma treating each interacting surface for at least 1 min, then incubating the assembled chip for at least 2 h at 80 °C. The morning of each experiment, bonded channels were cooled to room temperature and then treated with a 1:10 dilution of poly-L-lysine (Sigma catalog no. P8920) in milliQ water. Poly-L-lysine treatment increased the number of attached cells and extended attachment duration, allowing for longer observations of single cells without affecting growth (**Supplementary Fig. 2b, c**). This treatment has no effect on growth rate (**Supplementary Fig. 2c**) and involves incubating the diluted poly-lysine solution inside each channel for 15 min, before gently removing the solution and flushing the emptied channel with sterile milliQ water. The treated channels were then air dried for at least 2 h prior to experimental use.

### Nutrient signal calibration and generation

All fluctuating nutrient signals in this study switched between two nutrient media: a “high” and “low” concentration of LB. To switch between mediums, we oscillated pressure within each reservoir of nutrient medium while maintaining a steady mean pressure to ensure a steady total flow rate of medium through the device (**Supplementary Fig. 1**). This flow rate was determined by collecting the fluid output from the MSG and measuring the volume per minute. While the pressure differentials across the set-up can vary (*e.g*., different device, slight variations in tubing lengths and angles), the range of flow rates used had no effect on growth rate (**Supplementary Fig. 2**). Because the pressure differentials could vary from day-to-day, the pressure differences required to completely switch between mediums were calibrated prior to each experiment at 20× magnification, which enabled visualization of the entire signal junction (**Supplementary Fig. 1**). These calibrated pressure differences were then used to define the fluctuating nutrient signal. The pressure system is programmed to generate the signal in synchrony with image acquisition. Thus, timestamps from the image data can be directly correlate with specific time points within the nutrient signal. Separately, the stability of the calibrated signals was visually confirmed by comparing the fluorescent signal exiting the junction with the signal observed downstream (**Fig. 1c**; **Supplementary Fig. 3**). These visualizations were conducted at 60× magnification and quantified by custom image processing scripts in MATLAB (see **Data and Software Availability**).

### Quantification of switching timescale and nutrient signal stability

To assess the accuracy between our designed signal (an even square-wave) and the signal realized within the microfluidic device, we compared the transition dynamics of nutrient shifts occurring immediately after the signal junction and those occurring near the end of the cell imaging region. Two positions in the MSG were imaged while the fluid flowing through fluctuated between a medium labeled with 0.26 nM sodium fluorescein and an unlabeled medium on a 30 s period. Specifically, we compared the sharpness of the signal immediately upon generation with that observed further downstream, as experienced by the surface-attached cells and observed virtually no decay in the fluorescent signal between the signal junction and the imaging region (**Supplementary Fig. 3**), only the time delay as calculated in **Supplementary Table 1**. While no fluid mixing occurs in this device – we operate under laminar flow regimes and there is no Lagrangian mixing – the diffusion of nutrients (or sodium fluorescein, which is 2–4 times the molecular weight of an amino acid) could potentially smooth out our nutrient signal. Diffusion can be further aided by Taylor dispersion, a phenomenon in which velocity gradients in the fluid flow (*i.e*., shear) work to increase the effective diffusion of a chemical species by spreading it across a larger region, thereby favoring diffusion. Were the magnitude of these effects non-negligible in our system, we should expect different slopes during transitions between the signals at the junction and downstream. Specifically, the downstream signal should have a longer transition time as we would detect fluorescence leaching into the *C*_low_ phases of the signal. However, the time required to complete transitions (*i.e*., time to go from baseline to saturated fluorescent signal and vice versa) was about 2 s in both locations (**Supplementary Fig. 3**). Thus, our flow rates are sufficiently fast to carry our intended signal across the entire length of the device without noticeable smoothing from diffusion. We also determined that the periodic oscillations in nutrient signal are robust across time. The programmed period (*T* = 30 s) was reliably quantified between peaks and between troughs from the repetitive fluorescein signal (**Supplementary Fig. 3**).

### Microfluidics experimental procedure

The complete system involves: (1) a Nikon Eclipse Ti inverted microscope, (2) a full-case incubator that maintains a stable temperature (37 °C) around the entire microscope, except for the camera and light sources, (3) a computer to operate the microscope software (Nikon Elements) and MATLAB, (4) a data acquisition (DAQ) system that interfaces with MATLAB to control two pressure regulators, one for each nutrient source, (5) two reservoirs of nutrient medium, one of each nutrient concentration, and (6) a source of compressed air. The compressed air is fed into the pressure system through a manual regulator, which caps the pressure directed towards the two automated regulators at 1.5 psi. To ensure that the automated regulators receive a stable input, the pressure of the compressed air source is higher than this maximum value. Each automated regulator is connected to and modulates the internal pressure of one reservoir of nutrient medium (*C*_high_ or *C*_low_). Each nutrient reservoir is a septum-capped glass vial (vials: VWR, Cat. No. 548-0154; caps: VWR, Cat. No. 548-0872) with two needles inserted into the silica septum: one short and one long. The short needle directly connects an automated pressure regulator with the air space within its reservoir, thereby adjusting the pressure within the reservoir as dictated by the MATLAB signal (**Supplementary Fig. 1a**). The long needle connects the fluid within its reservoir with the microchannel via tubing inserted into the inlets of the device (**Supplementary Fig. 1a**). The microscope and media are contained within a custom LIS incubator, which maintains the sample and all media at 37 °C.

Experiments were based on the exposure of cells attached to the lower surface of the microchannels to precisely controlled fluctuating or steady nutrient conditions, and the imaging of thousands of cells in the downstream imaging region in order to calculate their individual growth rates. The treated, dry microchannels were inoculated with around 50 μl of the seed culture (see Culture procedure) for 10–15 min, allowing cells to settle and attach to the glass surface within each microchannel before flow was established. Prior to inoculation, the microfluidic device was placed in a vacuum for at least 10 min to remove air from the PDMS. This step helped to avoid the presence of bubbles inside the channels, by removing air from the PDMS so that any air introduced in the set-up would be absorbed by the PDMS. Inputs to fluctuating conditions were two septum-capped glass vials (one each for high and low nutrient) from which flow was driven by a custom-built air pressure system (**Supplementary Fig. 1a**). Inputs to steady conditions were 10 mL plastic syringes (Codan) from which flow was driven by a syringe pump (Harvard Apparatus). Outputs for all conditions led to liquid waste receptacles. To avoid changes in pressure throughout experiments, we ensured that the waste tubing was sufficiently short to never become submerged by the rising level of waste water.

### Image acquisition

Individual cells from all microchannel environments were imaged with phase contrast microscopy using a Nikon Eclipse Ti inverted microscope equipped with an Andor Zyla sCMOS camera (6.5 μm per pixel) at 60× magnification (40× objective with 1.5× amplification), for a final image resolution of 0.1083 μm per pixel. This magnification was high enough to detect changes in growth between each image, yet low enough to image hundreds of cells per field of view. Each position was repeatedly imaged every 117 s (1:57 min), a time step sufficiently high to allow the acquisition of multiple time points along a growth curve (*i.e*., 10 time points in a 20 min cell cycle), yet infrequent enough to image a total of 40–50 positions within each time step. Generally, 10 imaging positions per condition allowed us to track 500–1000 or more cells per nutrient condition. We confirmed that growth rates were independent of a cell’s position along the 10-mm-long region imaged within the microchannel (**Supplementary Fig. 4**) and therefore that cells experienced identical nutrient time series, regardless of location within the microchannel. The uneven time step (1:57 min as opposed to 2:00 min) was chosen to avoid potential aliasing effects, by sampling at various points along the nutrient period instead of repeatedly at the same few. Light exposure was limited to 20 ms per image, with the shutter only open during image capture. Image acquisition was fully automated through Nikon Elements, supplemented with the Nikon Perfect Focus System to prevent loss of focus due to vertical shifts in the sample.

### Image processing

In preparation for analysis, image sequences from microfluidic experiments were first passed through a particle tracking step and a quality control step. First, a custom MATLAB particle tracking pipeline was developed to: (1) read image data directly from Nikon Elements image files, (2) identify particles based on pixel intensity, (3) fit an ellipse to each particle and measure particle parameters (*e.g*., length, width) and (4) track individual particles through time. Second, to exclude errors from our analysis – for example, particles arising from noise (*i.e*., non-uniformity in the background) or particles that include more than one cell – a quality control step trimmed our tracked dataset, using size criteria and noise filters to exclude errors. The parameter values used in both steps ensured that the vast majority of tracks derive from isolated single cells. The final output of these two steps is a data matrix containing parameter data (*e.g*., cell length) over time for hundreds of individual cells growing in isolation. Reducing our analysis to cells without neighbors allowed us to assume no accumulation or depletion of medium components, as well as physical interactions between cells. Cells in contact were excluded from analysis to avoid the possibility of metabolic interactions and imprecision in the measurement of cell size. Specifics regarding the particle tracking and quality control steps are available in the scripts (see **Data and Software Availability**).

### Quantification and statistical analyses

#### Calculating instantaneous growth rate, μ

One widely used method to calculate growth rate is to consider single-cell growth an exponential process (11; 12) and solve for instantaneous growth rate, μ. This definition of growth rate is used throughout this study. From the length and width measured during particle tracking, the instantaneous volume of each individual cell was approximated as a cylinder with hemispherical caps (17). The approximated volumes were then used to compute instantaneous single-cell growth rates in terms of volume doublings per hour. Using *V*(*t*+Δ*t*) = *V*(*t*) · 2^μΔ*t*^, we calculated μ between each pair of time steps, with Δ*t* = 117 s (imaging frame rate). Specifically, we took the natural logarithm of each volume trajectory and calculated the slope between each point. Dividing the slope by the natural log of 2 changes the base of the exponential from *e* to 2. Thus, μ represents the exponential rate at which volume doubles.

#### Accounting for day-to-day variability in growth rate

While steady-state growth rates were generally reproducible (**Supplementary Fig. 5b; Supplementary Table 2**), we found that growth rate measurements performed on the same day (*i.e*., same seed culture) were moderately correlated (**Supplementary Fig. 5c**). This correlation indicates that slight differences between the seed culture (which was different for each experiment) contributed the differences in growth rate measured from identical conditions between experiments. Thus, when comparing growth rate across conditions (*e.g*., *G*_fluc_ and *G*_ave_), we compared measurements performed on the same day before comparing between experimental replicates, calculating the fraction of *G*_ave_ represented by the measured *G*_fluc_ from that same experiment before calculating statistics (*i.e*., mean and standard deviation of *G*_fluc_/*G*_ave_) across experiments. We used this same approach when comparing *G*_fluc_ to *G*_J_, which was calculated from each experiment’s *G*_low_ and *G*_high_, and *G*_fluc_ to *G*_low_. The alternative approach for this comparison would be to calculate the mean and standard deviation between experimental replicates *before* calculating fractions (*e.g*., *G*_fluc_/*G*_ave_) and combining error. Numerically, this alternative approach yields very similar results.

## Data and Software Availability

The data that support this study are available from the corresponding authors upon request. Image processing, data analysis and plotting scripts are available on Github: https://github.com/jkimthu/growing-up

## Acknowledgements

We are grateful to Michael Laub, Katharina Ribbeck and members of the Stocker and Ackermann groups for discussions regarding this work. Sine Svenningsen, Russell Naisbit, Johannes Keegstra and Juanita Lara Gutierrez provided critical readings on the manuscript. We also thank Suckjoon Jun, Sarah Cox and JT Sauls for the NCM3722 strain of *E. coli*. This work was funded by the Gordon and Betty Moore Foundation through Investigator Award GBMF3783 (to R.S.) and Grant GBMF3801 (to R.S.), by the Simons Foundation through Grants 542395 (to R.S.) and 542379 (to M.A.) as part of the Principles of Microbial Ecosystems (PriME) Collaborative, and by the Swiss National Science Foundation under Grant No. 315230_176189 (to R.S.).

## Author Contributions

J.N., V.F., M.A. and R.S. designed the study; J.N. and V.F. designed the microfluidic system, pressure control system/software, and particle tracking software; J.N. performed the microfluidic experiments and analyzed the growth data; S.P. performed MS measurements and analyzed the metabolomics data; U.S. provided overview on metabolomics data generation and analysis; J.N., M.A., and R.S. wrote the paper; all authors reviewed the paper before submission.

