## Supplementary Information for "A distinct growth physiology enhances bacterial growth under rapid nutrient fluctuations"

**Supplementary Fig. 1: The Microfluidic Signal Generator (MSG) delivers fluctuating nutrient signals alongside steady ones.**

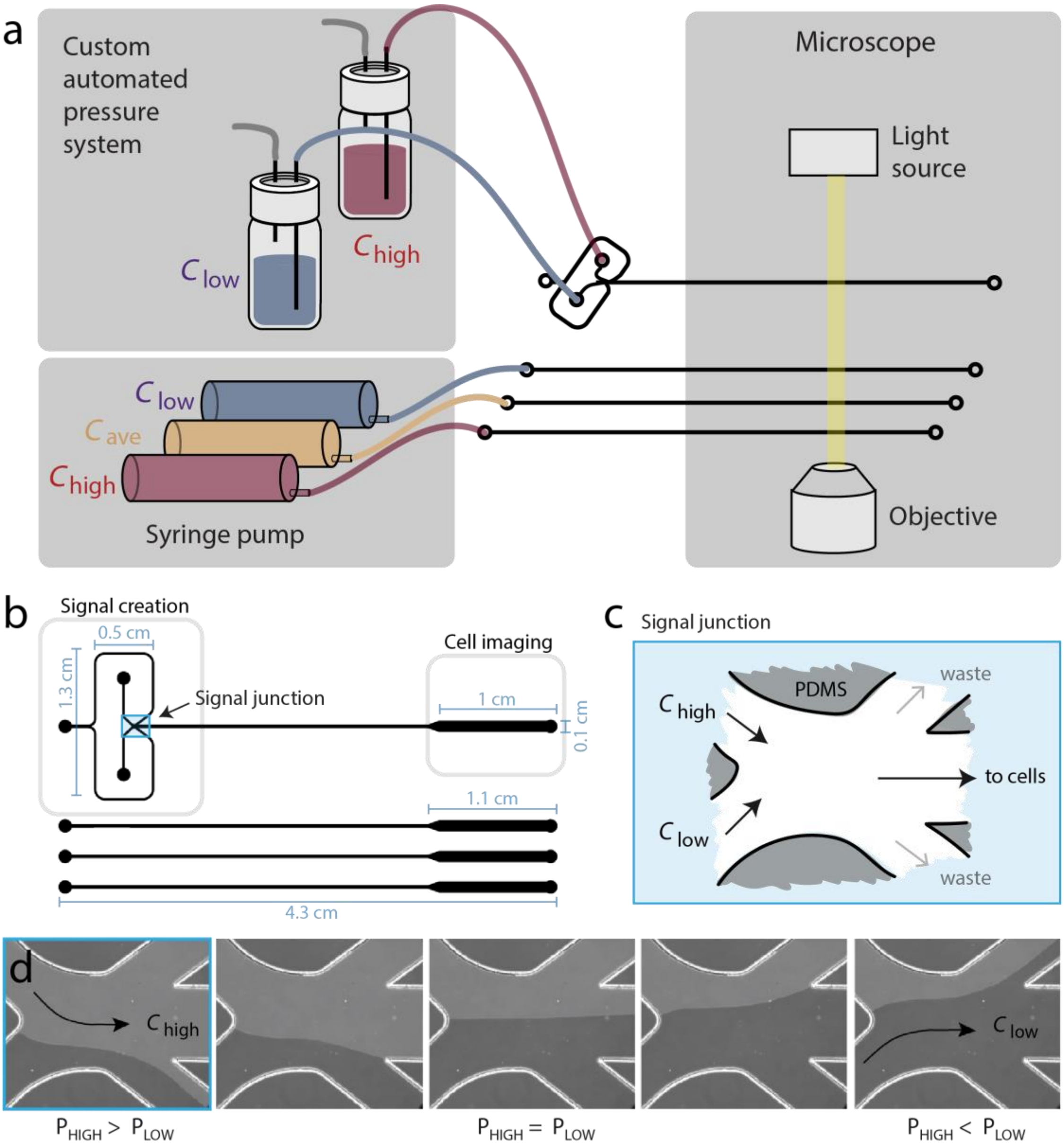


**a** Four nutrient signals are delivered within each experiment, in parallel microfluidic channels imaged by the microscope. The fluctuating signal is produced by oscillating the pressure in two media vials, one containing the high nutrient concentration ($C_{high}$) and one containing the low concentration ($C_{low}$), at a user-programmed frequency. The programmed signal automates which vial has the higher internal pressure, and the system is calibrated such that, when the pressure in the $C_{high}$ vial is higher, only $C_{high}$ reaches the cells downstream. The fluid input to each steady environment is a single syringe filled with either $C_{low}$, $C_{ave}$ or $C_{high}$. All three syringes are pushed by the same syringe pump at a flow rate of 15 µL/min. In the diagram, the colored lines represent the flexible polyethylene tubing connecting each fluid input with the microchannel inlets. The circular microchannel features not associated with inputs are the microchannel outlets, which are connected via tubing to waste receptacles (not shown). **b** 2-D design of microfluidic device. The channel height (not shown) is uniformly 60 µm. The top channel with the unique upstream feature is the MSG, and the three straight channels deliver the control steady environments. In all channels, the nutrient signal travels over 2.5 cm before reaching the cells. **c** The signal junction upstream in the MSG has two inlets and three outlets: one inlet per medium ($C_{low}$ and $C_{high}$), one outlet towards the cells, and two waste outlets to remove excess fluid that does not enter the middle channel. **d** Nutrient switches are achieved via pressure oscillations in the switching junction. When the pressure in the low nutrient reservoir is approximately equal to that in the high nutrient reservoir, then both media flow to the cells downstream. To generate square waves that oscillate between $C_{low}$ and $C_{high}$, we calibrated the pressure ratios required such that only $C_{low}$ or only $C_{high}$ would enter the middle outlet toward the cells.

**Supplementary Fig. 2: Nutrient concentration, not fluorescein, flow rate nor poly-L-lysine treatment, controls growth rate.**

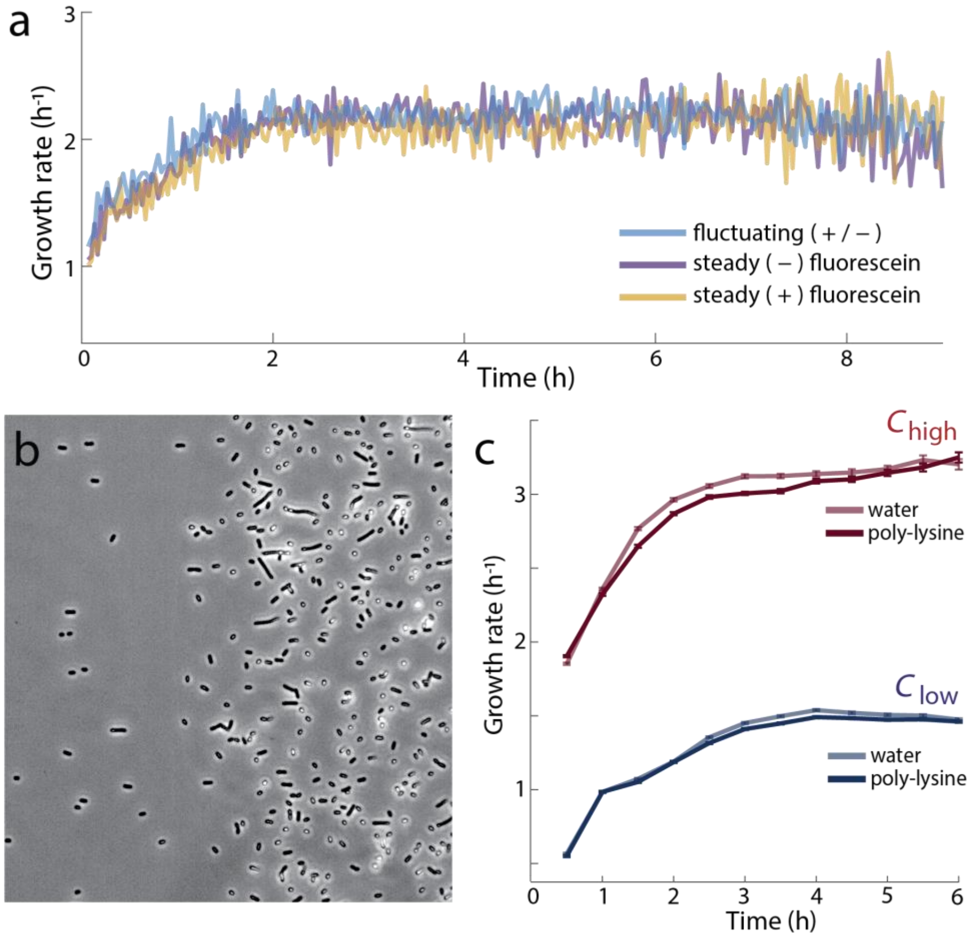


**a** Neither fluorescein labeling nor flow rate affect growth rate. Time-evolution of growth rate in three parallel channels: (1) fluctuations between fluorescein-labeled (0.26 nM) and unlabeled nutrient on a 30 s period, (2) steady nutrient without fluorescein, and (3) steady nutrient with 0.26 nM fluorescein. All nutrient media are 1% LB diluted in equimolar salt solution, approximately equal to $C_{ave}$ (1.05% LB). Flow rate within the fluctuating channel is 22 µL min$^{-1}$, as generated by the compressed air-based pressure system. Flow rate within the steady channels is 15 µL min$^{-1}$ generated by a syringe pump. Each curve represents the mean instantaneous growth rate for each condition. **b** Poly-L-lysine treatment enhances bacterial attachment. A phase contrast image of a microchannel with *E. coli* attached to the lower, glass surface in flow. The right-hand side of the image was treated with poly-lysine prior to inoculation while the left-hand side was not. The boundary of the poly-lysine treatment is clearly marked by the increased presence of adherent cells. **c** Poly-L-lysine treatment does not affect growth rate. Mean instantaneous growth rate over time under steady nutrient conditions demonstrates that cells surface-attached to a poly-L-lysine-treated surface display no difference in growth rate compared to cells surface-attached to an untreated glass surface. Two channels delivered $C_{high}$ and two channels delivered $C_{low}$, of which one of each was treated with poly-L-lysine and the other with sterile milliQ water.

**Supplementary Fig. 3: Media switching occurs within 3 s at user-defined frequencies.**

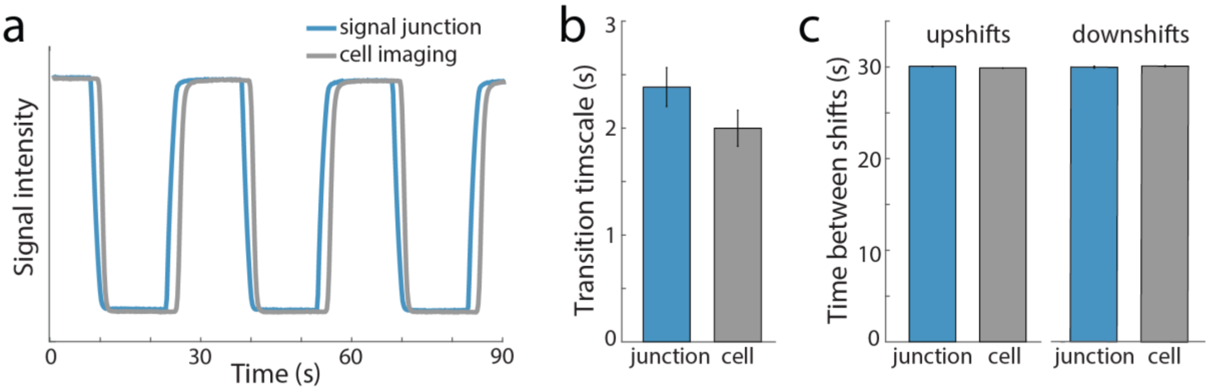


**a** Fluctuations between two media, one of which was labeled with 0.26 nM sodium fluorescein, at the signal junction and near the end of the cell-imaging region of the MSG. A switch is considered complete when the fluorescence intensity reaches the baseline or the saturated intensity. **b** Measured transition time at the signal junction and the end of the cell-imaging region. Transition time is defined as the time required to fully switch between media. Error bars are standard deviation from six transitions. **c** Period lengths measured from the fluorescent signal, which was programmed to oscillate on a 30 s period, demonstrate that the periodic oscillations are robust. Upshifts measure the time between peak fluorescent signals. Downshifts measure the time between troughs. Error bars represent standard deviations between three transitions. The programmed period was reliably quantified between peaks and between troughs from the repetitive fluorescein signal.

**Supplementary Fig. 4: Growth rate does not vary with distance from the signal junction**.

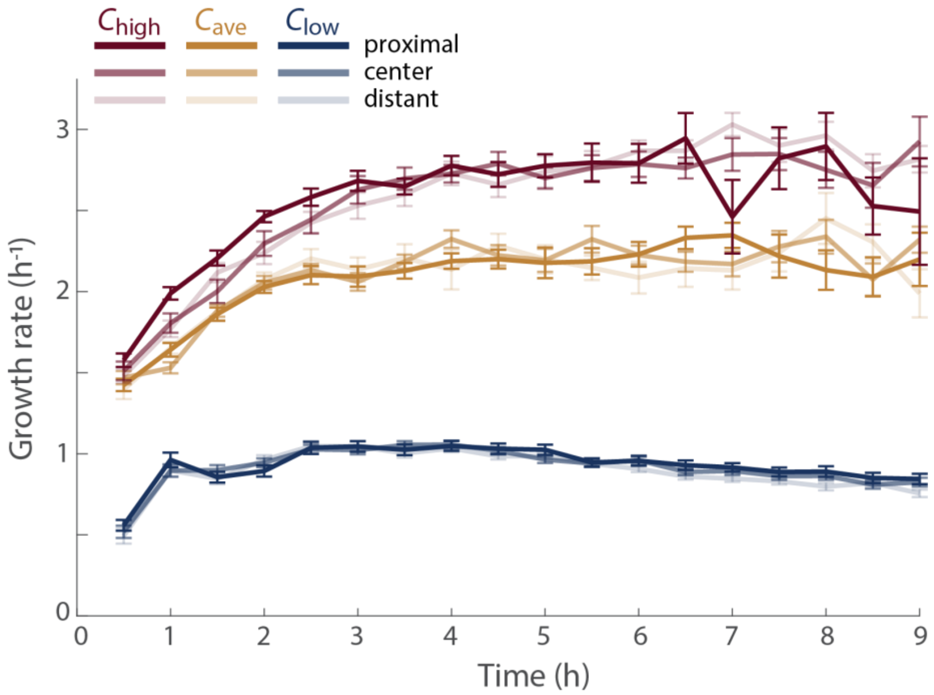


Mean instantaneous growth rate over time from three parallel nutrient concentrations: $C_{low}$ (0.1% LB, blue), $C_{ave}$ (1.05% LB, yellow) and $C_{high}$ (2% LB, red). Cells growing in each concentration were imaged at 10 distinct imaging positions along the microchannel, three of which are shown here represented by lines of different transparency. Proximal refers to the position within the imaging region closest to the signal junction, whereas distant refers to the position furthest from the junction. The lack of systematic variation in growth rate with position suggests that, as expected from the volume of medium present and the flow rate, the nutrient medium was not depleted by cells growing within the microchannel.

**Supplementary Fig. 5: Steady-state growth rates under a range of LB dilutions.**

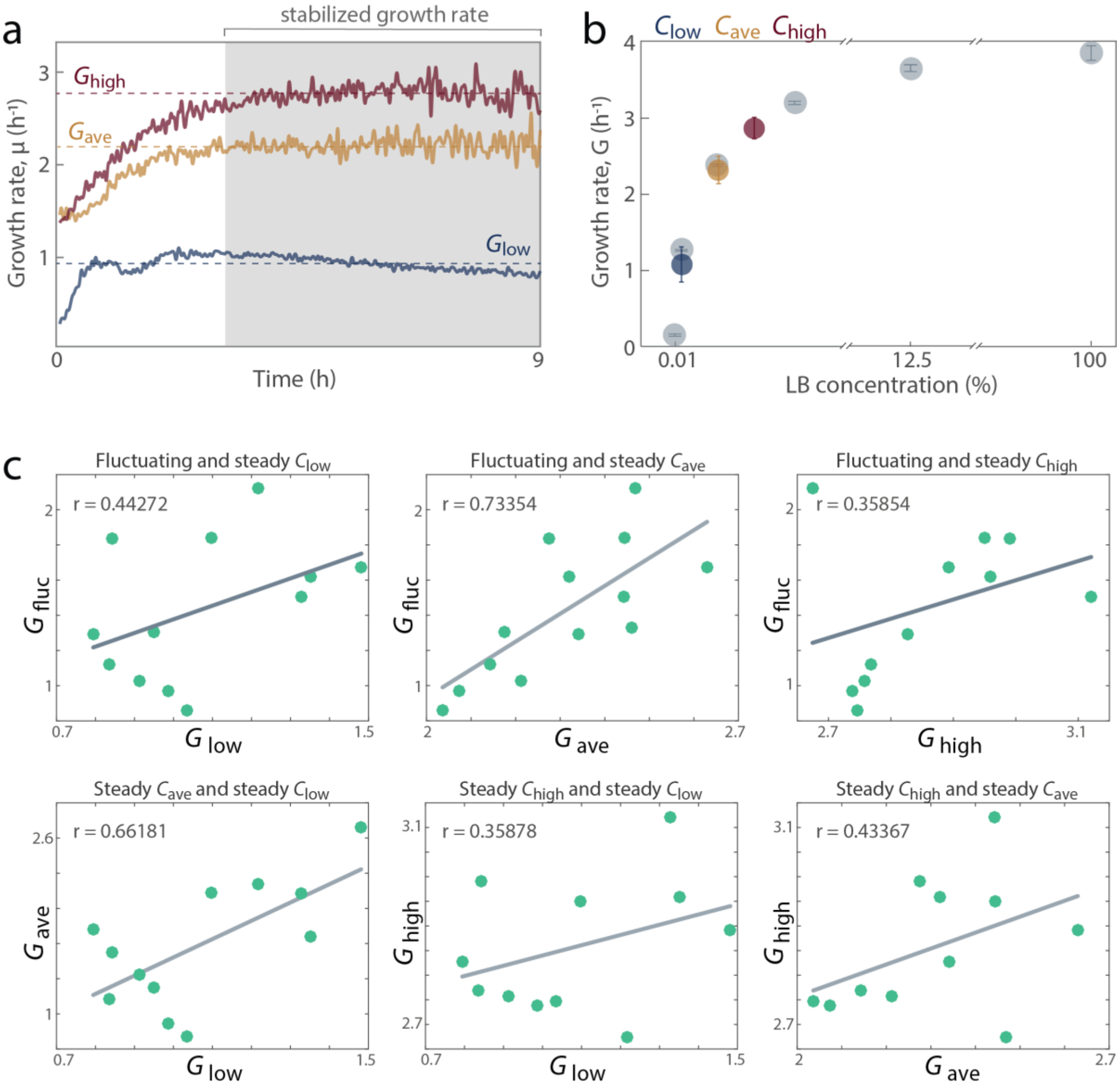


**a** Growth rate evolving over time in steady $C_{low}$, $C_{ave}$ and $C_{high}$. Stabilization of instantaneous growth rate was consistently achieved within the first 3 h of the onset of nutrient delivery. We defined the steady-state growth rate measured from a given condition as the time-average of all instantaneous growth rate data collected after this 3 h mark, indicated by the dashed line and annotated as $G_{low}$, $G_{ave}$ and $G_{high}$. **b** Growth rate within our device under various concentrations of LB was empirically characterized from six steady nutrient conditions, ranging from 0.01% to 100% LB. These six steady-state growth rates (gray) – 0.1446 ± 0.0057 (0.01% LB); 1.2622 ± 0.0042 (0.1% LB); 2.3748 ± 0.0057 (1% LB); 3.1924 ± 0.0086 (3.125% LB); 3.6411 ± 0.0170 (12.5% LB); 3.8443 ± 0.0960 (100% LB), all mean ± s.e.m. in units of $h^{-1}$ – were used to reconstruct a Monod curve from which the experimental concentrations were chosen. The final $C_{high}$ (2% LB, red), $C_{ave}$ (1.05% LB, yellow) and $C_{low}$ (0.1% LB, blue) nutrient conditions were chosen such that $G_{ave}$ was clearly distinguishable from $G_{low}$ and $G_{high}$. Color circles represent the mean steady-state growth rate measured from $C_{low}$, $C_{ave}$ and $C_{high}$ with error bars representing the standard deviation among 11–13 replicates (Supplementary Table 2). **c** Daily correlations between the time-averaged growth rate (*G*) of different nutrient conditions. Each point in the plot denotes the time-averaged growth rate of two conditions, measured from the same four-condition experiment (i.e., conducted on the same day, on the same microfluidic chip and seeded simultaneously with the same starting culture). The gray line is the fitted linear regression, and *r* is the corresponding correlation. These moderate correlations between conditions performed on the same day indicates that slight differences between the seed culture contributed variability in growth rate measured from identical conditions performed on different days. Thus, we compared growth rate across conditions performed on the same day before comparing experimental replicates.

**Supplementary Fig. 6: Metabolites are consumed proportionally in the studied nutrient regimes.**

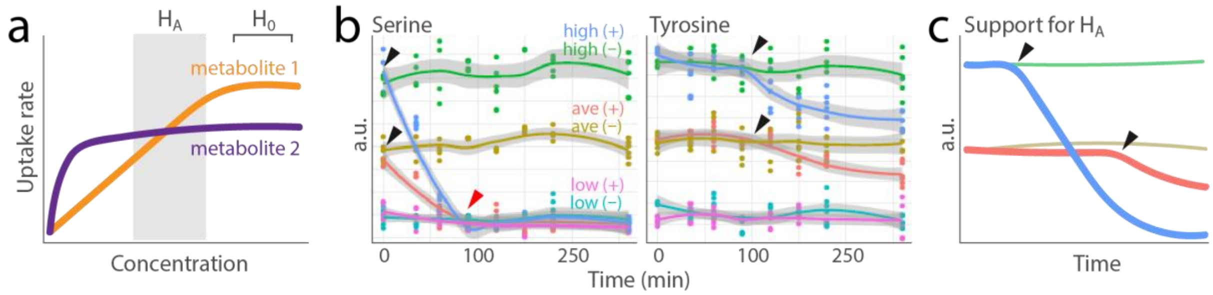


**a** Different nutrient uptake kinetics could potentially affect the proportional uptake between different substitutable metabolites. If $C_{low}$, $C_{ave}$ and $C_{high}$ were to span the regime shaded by the grey box, then metabolite 1 would be the primary nutrient source in $C_{high}$, whereas metabolite 2 would be the primary nutrient source in $C_{low}$. In this case, the difference in growth rate between $G_{low}$ and $G_{high}$ would support the hypothesis that differences in growth rate between nutrient concentrations derived from differences in the ratios of metabolites consumed ($H_A$). Were our chosen concentration regime to fall outside of the shaded region, then we would expect Monod growth in which the primary nutrient source is constant across LB dilutions and the change in growth with nutrient concentration results solely from change in the uptake rate of that nutrient ($H_0$). **b** Mass spectrometry detection of metabolic consumption supports $H_0$, because each nutrient species is used simultaneously across nutrient concentrations. In a fixed initial medium (no flow) inoculated with *E. coli* (+), serine is consumed (black arrows) until it can no longer be detected (red arrow), at which point tyrosine begins to decrease, in both $C_{ave}$ and $C_{high}$. The detection curves for these two metabolites are shown as representatives for the 284 analyzed metabolites (Supplementary Table 6). Each curve denotes the detected level of a specific metabolite in $C_{low}$, $C_{ave}$ and $C_{high}$ with (+) or without (-) the addition of growing cells at $t$ = 0. Each point indicates the value of one replicate and the line the mean across replicates ($n$ = at least 3 per time point). In all conditions, OD600 saturates within 300 min. Black arrows indicate where metabolite level begins to decrease. **c** Were $H_A$ to be true, we would expect the same nutrient species to be used at different time points across different nutrient concentrations, due to changes in the primary nutrient with variation in medium concentration. Among the 284 metabolites detected, we never observed a metabolite profile that supported $H_A$.

**Supplementary Fig. 7: Representative single-cell μ trajectories demonstrate that growth rate dynamics, while noisy, occur in individual cells.**

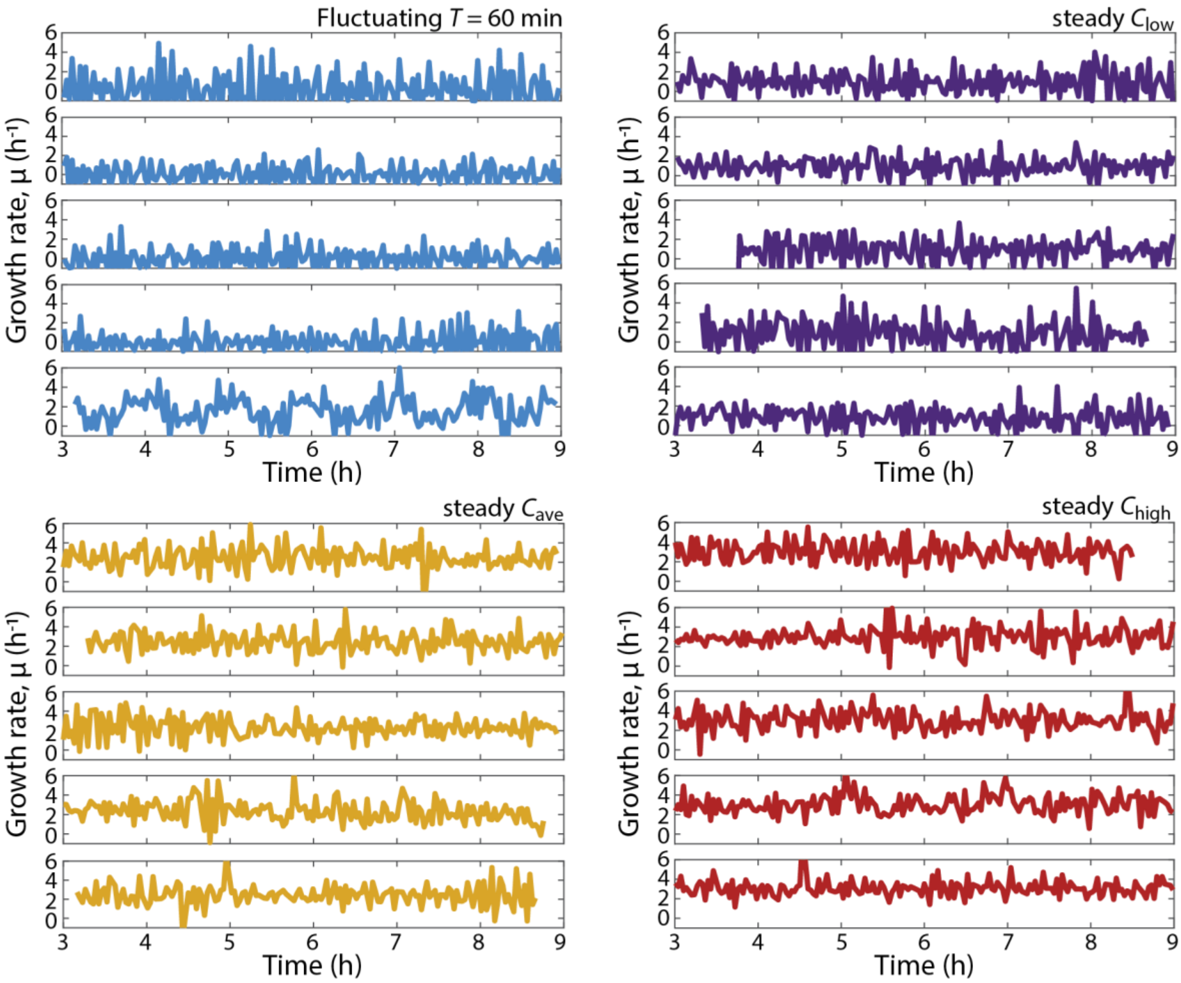


Visualized instantaneous growth rate dynamics from representative single cell lineages growing in either 60 min nutrient fluctuations (blue) or steady $C_{low}$ (purple), $C_{ave}$ (yellow) and $C_{high}$ (red). For each trajectory, each time step between instantaneous growth rates (μ) was 117 s, with the exception of time steps in which a cell division occurred (leading to a momentary strong negative growth rate), which were omitted for clarity. Due to the high variability in μ measured from time step to time step, growth rate responses to changes in nutrient conditions required averaging across several single cells to detect a significant signal. Still, periodic fluctuations in growth rate are occasionally were occasionally visible by eye from individual trajectories under fluctuating nutrient conditions (bottom most panel from *T* = 60 min).

**Supplementary Fig. 8: Frequency of nutrient shifts determine fraction of timesteps containing shifts and change in growth rate after nutrient upshift.**

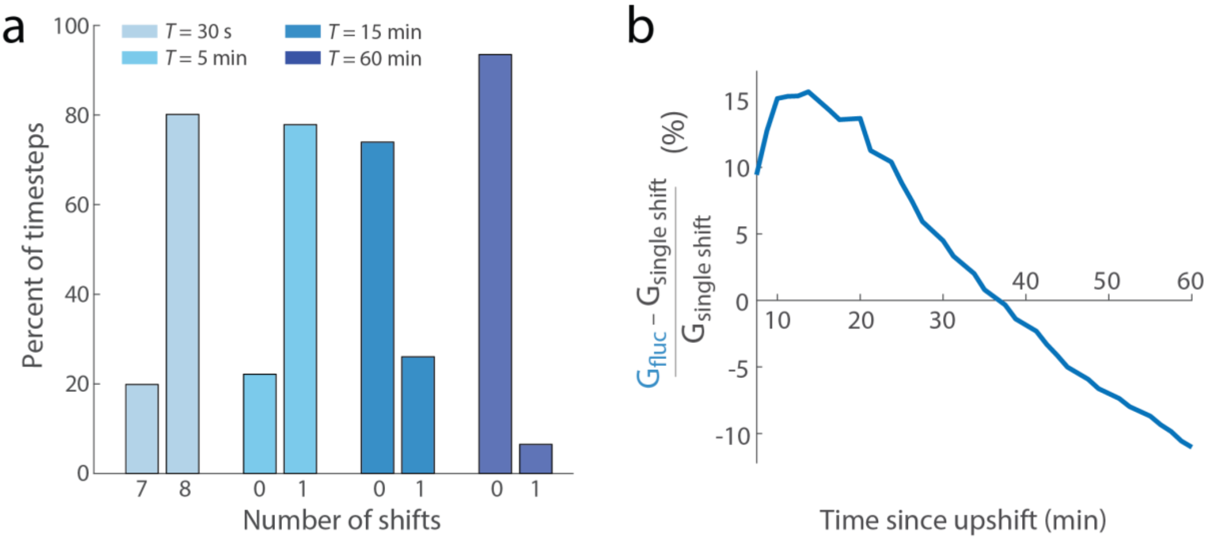


**a** Percentage of instantaneous growth rate measurements encompassing a nutrient shift for each nutrient signal timescale. A 10 h experiment imaged every 117 s comprises 307 imaging intervals. Given the periods of the nutrient signals delivered (30 s, 5 min, 15 min and 60 min), we calculated the percentage of imaging intervals for each nutrient signal that contained each possible number of nutrient shifts. Each imaging intervals in the fastest nutrient signal (30 s) contains either 7 or 8 nutrient shifts; imaging intervals in longer nutrient signals have either 0 or 1. The decreasing fraction of imaging intervals that contain a nutrient shift with increasing length of the nutrient signal is consistent with the decreasing strength of "anticipation", i.e., the phase shift by which growth rate appeared to respond to nutrient shifts before the shift occurred. This calculation is evidence that this phase shift in growth rate response, particularly evident in the 5 min and 15 min nutrient signals (Fig. 2c), resulted from the smoothing of the growth rate signal during analysis, not a biological response in the experiment. **b** Cells adapted to rapid nutrient fluctuations have a growth advantage in the first 30–40 minutes after a nutrient shift. The curve shows the relative difference between the growth rate of fluctuation-adapted cells (*T* = 60 min) and the growth rate of cells grown at steady-state that experience a single nutrient shift as a function of time since a nutrient upshift. The positive value in the first tens of minutes indicates a growth advantage for fluctuation-adapted cells after a nutrient shift. At 30 min after the shift, the growth rate in the fluctuating condition is considered stabilized and used to calculate percent differences from single upshift data extending beyond 30 min (Fig. 4a, b).

**Supplementary Fig. 9: Predicted growth rate dynamics under fluctuations based on the dynamics observed after single shifts.**

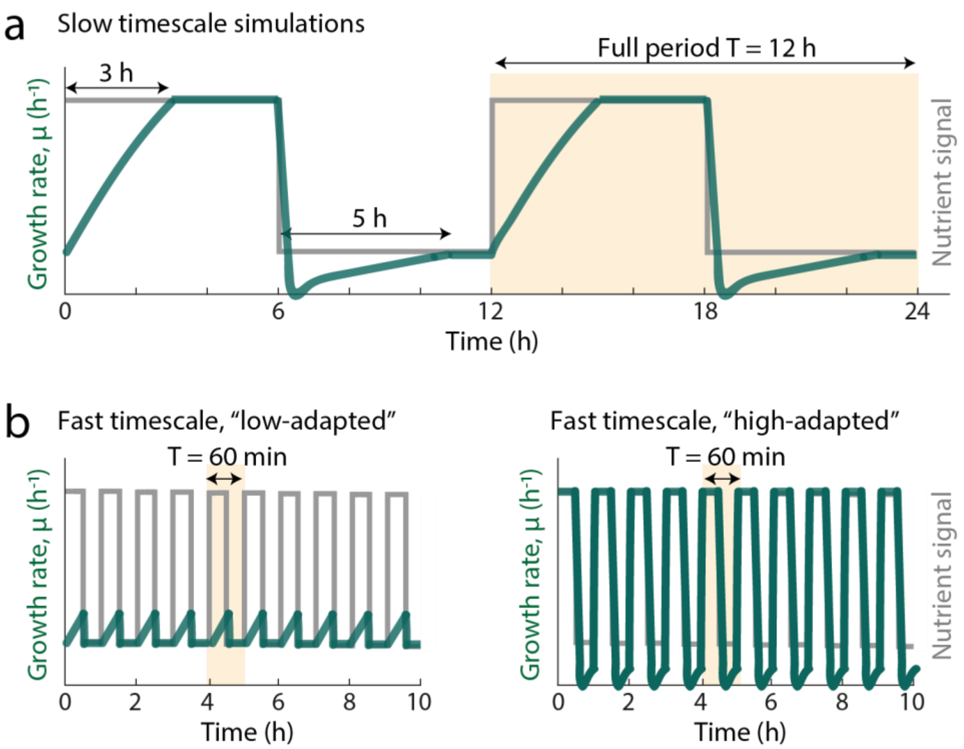


**a** At slow timescales ($T$ = 12 h and greater), each nutrient phase ($C_{high}$ or $C_{low}$) is longer than the stabilization time required to transition in physiology between steady states (3 h to reach $G_{high}$ after an upshift, 5 h to reach $G_{low}$ after a downshift). The growth rate dynamics (green) during these transitions is predicted based on the dynamics measured in single-shift experiments (**Fig. 4a**). The predicted $G_{fluc}$ value is simply a time-average of instantaneous growth rates for one full period. **b** At fast timescales (T = 60 min or less), each nutrient phase is shorter than the stabilization time required to transition between steady states, and our null models predicted lower values of $G_{fluc}$ for faster nutrient fluctuations. We considered two alternative scenarios. In the "low-adapted" simulations (left), time spent in $C_{low}$ is considered as growth at $G_{low}$, and the time in $C_{high}$ is modeled on the basis of the beginning of the physiological transition observed after a single upshift. In the "high-adapted" simulations (right), time spent in $C_{high}$ is considered as growth at $G_{high}$, and the time in $C_{low}$ is modeled on the basis of the beginning of the physiological transition observed after a single downshift. The "high-adapted" simulation accommodates the likelihood that repeated exposure to a nutrient concentration (*i.e., $C_{high}$*) can prepare cells for steady state growth in that concentration (*i.e., $G_{high}$*). Thus, the "high-adapted" simulation serves as a generous upper bound for our predictions. All simulated $G_{fluc}$ are displayed in **Supplementary Table 5**.

**Supplementary Table 1**

**Mean lag times between signal generation and signal delivery to cells, per experiment.** Lag time was calculated from the flow rate and effective cross-sectional area. Error denotes the standard deviation of the 10 cell positions sampled per experiment, across the length of the imaging region. Each straight channel delivers fluid from one steady source, a 10 mL syringe loaded into the same Harvard Apparatus syringe pump, which pushes the fluid into the channels at a flow rate of 15 µL $min^{-1}$. The different flow rates used (15–27 µL $min^{-1}$) did not affect growth rate (Supplementary Fig. 2a).

| Experimental condition | Flow rate, *Q* (µL $min^{-1}$) | Lag time (s) |
|---|---|---|
| 30 s period (2017-11-12) | 15 | 2.94 ± 0.12 |
| 30 s period (2017-11-14) | 21 | 2.08 ± 0.08 |
| 30 s period (2018-01-04) | 27 | 1.63 ± 0.06 |
| 5 min period (2017-10-10) | Not recorded | N.A. (assigned lag time = 1.7 s) |
| 5 min period (2017-11-15) | 21 | 2.08 ± 0.08 |
| 5 min period (2018-01-11) | 21 | 2.07 ± 0.07 |
| 15 min period (2017-11-13) | 20 | 2.20 ± 0.08 |
| 15 min period (2018-01-12) | 22 | 1.97 ± 0.07 |
| 15 min period (2018-01-16) | 22 | 1.98 ± 0.07 |
| 15 min period (2018-01-17) | 23 | 1.89 ± 0.07 |
| 60 min period (2018-01-29) | 26 | 1.67 ± 0.05 |
| 60 min period (2018-01-31) | 26 | 1.69 ± 0.07 |
| 60 min period (2018-02-01) | 26 | 1.70 ± 0.07 |

**Supplementary Table 2**

**Summary table of steady-state growth rates.** Mean growth rate (*G*) and standard error of the mean for all conditions from all replicate fluctuating experiments. Cells labelled with N.A. represent instances in which a bubble formed within the channel during the first 3 h of data acquisition, preventing the determination of a steady-state growth rate. Units of growth rate are $h^{-1}$. The column labelled "Experiment" lists the date on which the experiment was performed. Each row associated with one fluctuation timescale represents a distinct experimental replicate carried out simultaneously in one microfluidic device.

| | Experiment | Fluctuating | Steady low | Steady average | Steady high |
|---|---|---|---|---|---|
| 30 s | 2017-11-12 | 1.8391 ± 0.0040 | 1.0965 ± 0.0024 | 2.4439 ± 0.0053 | 2.9497 ± 0.0067 |
| | 2017-11-14 | 2.1229 ± 0.0040 | 1.2170 ± 0.0038 | 2.4692 ± 0.0087 | 2.6744 ± 0.0183 |
| | 2018-01-04 | 1.8373 ± 0.0077 | 0.8430 ± 0.0036 | 2.2755 ± 0.0092 | 2.9907 ± 0.0109 |
| 5 min | 2017-10-10 | 1.6728 ± 0.0063 | 1.4814 ± 0.0050 | 2.6296 ± 0.0105 | 2.8919 ± 0.0143 |
| | 2017-11-15 | 1.6198 ± 0.0075 | 1.3525 ± 0.0054 | 2.3198 ± 0.0098 | 2.9588 ± 0.0110 |
| | 2018-01-11 | 1.3032 ± 0.0037 | 0.9503 ± 0.0018 | 2.1744 ± 0.0088 | N.A. |
| 15 min | 2017-11-13 | 1.5064 ± 0.0109 | 1.3282 ± 0.0061 | 2.4434 ± 0.0105 | 3.1206 ± 0.0150 |
| | 2018-01-12 | 1.3302 ± 0.0077 | N.A. | 2.4605 ± 0.0083 | N.A. |
| | 2018-01-16 | 0.9695 ± 0.0123 | 0.9870 ± 0.0024 | 2.0733 ± 0.0151 | 2.7382 ± 0.0156 |
| | 2018-01-17 | 0.8595 ± 0.0123 | 1.0341 ± 0.0029 | 2.0362 ± 0.0141 | 2.7467 ± 0.0092 |
| 60 min | 2018-01-29 | 1.2941 ± 0.0096 | 0.7944 ± 0.0026 | 2.3412 ± 0.0055 | 2.8273 ± 0.0050 |
| | 2018-01-31 | 1.1211 ± 0.0129 | 0.8357 ± 0.0026 | 2.1422 ± 0.0063 | 2.7684 ± 0.0096 |
| | 2018-02-01 | 1.0272 ± 0.0091 | 0.9130 ± 0.0022 | 2.2123 ± 0.0076 | 2.7572 ± 0.0074 |

**Supplementary Table 2**

**Summary table of steady-state growth rates.** Mean G of replicates for each condition, reported with standard deviation (st dev), standard error of the mean (s.e.m.) and number of experimental replicates (n). For fluctuating conditions, the number in the parentheses represents the timescale of nutrient fluctuation in minutes, except for when noted (30 s).

| | $G_{low}$ | $G_{ave}$ | $G_{high}$ | $G_{fluc}$ (30 s) | $G_{fluc}$ (5) | $G_{fluc}$ (15) | $G_{fluc}$ (60) |
|---|---|---|---|---|---|---|---|
| mean: | 1.0694 | 2.3094 | 2.8567 | 1.9331 | 1.5319 | 1.1664 | 1.1474 |
| st dev: | 0.2271 | 0.1768 | 0.1363 | 0.1644 | 0.1998 | 0.3030 | 0.1354 |
| s.e.m.: | 0.0656 | 0.0490 | 0.0411 | 0.0949 | 0.1154 | 0.1515 | 0.0782 |
| n: | 12 | 13 | 11 | 3 | 3 | 4 | 3 |

**Supplementary Table 3**

**Predicted differences in biovolume production across fluctuation timescales, relative to the steady average environment**. The mean growth rate measured from each fluctuating environment, $G_{fluc}$, was used to calculate the daily biovolume (daily $M(t)$) expected to be generated per 1 $\mu m^3$ cell. Relative to the daily biovolume expected to be produced at the mean growth rate under the steady average conditions, $G_{ave}$, cells in the fluctuating environments are predicted to produce $10^2$–$10^8$ times less biovolume per day.

| Growth condition (*G*) | Mean *G* ($h^{-1}$) | Predicted daily *M(t)* | Fold difference, $M(t)/M(t)_{ave}$ |
|---|---|---|---|
| Steady average ($G_{ave}$) | 2.31 | $5 \times 10^{16}$ | 1 |
| T = 30 s ($G_{fluc,30}$) | 1.93 | $9 \times 10^{13}$ | $5 \times 10^{2}$ |
| T = 5 min ($G_{fluc,5}$) | 1.53 | $1 \times 10^{11}$ | $5 \times 10^{5}$ |
| T = 15 min ($G_{fluc,15}$) | 1.15 | $2 \times 10^{8}$ | $2 \times 10^{8}$ |
| T = 60 min ($G_{fluc,60}$) | 1.15 | $2 \times 10^{8}$ | $2 \times 10^{8}$ |

**Supplementary Table 4**

**Stabilized growth rate in fluctuations as a percentage of steady-state growth rate.** Under fluctuating nutrient environments, growth rates stabilize within 2–3 min of a nutrient shift. The second column quantifies the mean stabilized growth rate and standard deviation between replicates (*n* = 3 or 4). These stabilized growth rates are considerably lower than the steady-state rate corresponding to the post-shift environment: $G_{high}$ (2.86 ± 0.14 $h^{-1}$) in the case of upshifts, $G_{low}$ (1.07 ± 0.23 $h^{-1}$) in the case of downshifts. The reduction in growth rate relative to the "target" steady-state is quantified in column three (% loss = $\frac{G_{steady}-G_{fluc}}{G_{steady}} \cdot 100$).

| | Fluctuation timescale | Stabilized growth rate ($h^{-1}$) | % loss from steady-state |
|---|---|---|---|
| Upshift | 15 min | 1.86 ± 0.47 | 35.0 ± 0.2 |
| | 60 min | 1.86 ± 0.13 | 35.0 ± 0.1 |
| Downshift | 15 min | 0.65 ± 0.22 | 39.3 ± 0.3 |
| | 60min | 0.60 ± 0.10 | 43.9 ± 0.3 |

**Supplementary Table 5**

**Predictions of $G_{fluc}$ based on the growth rate dynamics observed after single nutrient shifts.** Values are calculated based on measured single-shift or steady-state data by one of three methods: slow for long timescales of nutrient shifts ($T$ = 12 h and greater), and low-adapted or high-adapted for faster nutrient timescales (Supplementary Figure 9). All $G_{fluc}$ are reported as a fraction of $G_{ave}$ (2.31 $h^{-1}$).

| Timescale ($T$) | Slow | Low-adapted | High-adapted |
|---|---|---|---|
| 96 h | 0.830 | N.A. | N.A. |
| 48 h | 0.810 | N.A. | N.A. |
| 24 h | 0.769 | N.A. | N.A. |
| 12 h | 0.688 | N.A. | N.A. |
| 60 min | N.A. | 0.563 | 0.644 |
| 15 min | N.A. | 0.511 | 0.638 |
| 5 min | N.A. | 0.502 | 0.623 |
| 30 s | N.A. | 0.497 | 0.619 |

**Supplementary Table 6**

**Annotations and masses of the 284 detected metabolites.** The table gives the Kegg annotations of all metabolites detected from the flask cultures of $C_{low}$, $C_{ave}$ and $C_{high}$ from time-of-flight mass spectrometry. These metabolites represent the metabolites from the media not yet consumed by the cultures as well as metabolites secreted into the media by cells. The ion intensity measurements from all 284 detected metabolites across all time points and replicates is available as an Excel file titled "metabolomics_intensities.xlsx" on GitHub: https://github.com/jkimthu/growing-up.
